# The mechanistic basis of cargo selection during Golgi maturation

**DOI:** 10.1101/2025.06.26.661719

**Authors:** Rebecca J Taylor, Nikita Zubkov, Katarzyna A Ciazynska, Jonathan GG Kaufman, Grigory Tagiltsev, David J Owen, John AG Briggs, Sean Munro

## Abstract

The multiple cisternae of the Golgi apparatus contain resident membrane proteins crucial for lipid and protein glycosylation. How Golgi residents remain in their designated compartments despite a constant flow of secretory cargo is incompletely understood. Here, we determined the structure of the COPI vesicle coat containing GOLPH3, an adaptor protein that binds the cytosolic tails of many Golgi residents. The structure, together with structure-guided mutagenesis and functional assays, reveals how GOLPH3 uses coincidence detection of COPI and lipid to engage Golgi residents preferentially at late cisternae. Our findings rationalize the logic of cisternal maturation and explain how COPI can engage different types of substrates in different Golgi cisternae to retrieve some proteins back to the ER while retaining others within the Golgi apparatus.

## Main Text

The Golgi apparatus is the central sorting station in the secretory pathway, with newly made lipids and secretory and membrane proteins arriving from the endoplasmic reticulum (ER), passing through the stack of Golgi cisternae and then being routed into the secretory or endocytic systems (*1*, *2*). A fleet of enzymes resident in the Golgi modifies the lipids and proteins passing through the stack. These enzymes include the many glycosyltransferases that generate the diversity of glycan structures found on glycolipids and glycoproteins and which underlie numerous biological processes (*3*, *4*).

Traffic between cellular compartments is mediated by carriers that are formed by coat proteins which select specific cargoes and then deliver them to the correct destination. Thus COPII-coated vesicles deliver cargo from the ER to the cis Golgi, while COPI-coated vesicles recycle escaped ER proteins back to the ER from the cis Golgi (*5*, *6*). The mechanism of transport through the Golgi stack has been proposed to occur either by vesicles moving cargo forward between cisternae, or by the cisternae maturing through the stack with vesicles recycling Golgi resident proteins back to earlier cisternae to maintain their distribution (*7*, *8*). COPI is the only vesicle coat found on the Golgi stack where it is present on coated-pits and vesicles from multiple cisternae (*9*, *10*). Thus, if the cisternal maturation model is correct then COPI would also be responsible for recycling Golgi residents within the stack. This raises the question of how COPI can recruit Golgi residents only in later cisternae and hence avoid transporting them back to the ER with the escaped ER residents it recruits at the cis Golgi.

COPI is formed by a heteromeric heptamer called coatomer that is recruited to membranes by the GTPase Arf1 (*10*, *11*). COPI has been shown to recycle escaped ER membrane proteins by binding directly to retrieval signals in their cytoplasmic tails such as K(x)Kxx (where x is any amino acid). These motifs bind to the membrane-proximal beta-propellor domains of α-COP and β’-COP (*12*, *13*). How Golgi resident proteins are incorporated into COPI vesicles is less clear. The resident glycosyltransferases of the Golgi are typically type II membrane proteins with a short cytoplasmic tail, a single transmembrane domain (TMD), and a luminal catalytic domain. For several such enzymes the cytoplasmic tails have been found to confer Golgi retention, with the TMD also contributing in some cases (*14*, *15*). Studies in yeast showed that a cytosolic protein Vps74 can bind the tails of glycosyltransferases and is required for their retention, and that it can also bind to the Golgi-localised lipid PI4P and to COPI through an unstructured N-terminal domain (*16*, *17*). Similar findings have been made for the mammalian Vps74 orthologue GOLPH3 (*18–21*). These observations suggest that Vps74 and GOLPH3 serve as adaptors to direct the recruitment of glycosyltransferases into COPI-coated vesicles. Deletion of GOLPH3 and its closely related paralogue GOLPH3L from cell lines results in extensive defects in glycosylation and GOLPH3 is found to be frequently amplified in some solid tumour types, consistent with the known role of glycans in regulating immune responses to tumour cells (*22–24*). Where GOLPH3 binds to COPI and how it recognises the tails of glycosyltransferases remain unknown. Several motifs for GOLPH3 binding have been suggested, typically comprised of combinations of basic residues and, in some cases, hydrophobic ones (*14*, *15*, *20*, *25*, *26*).

Here we report a structure of COPI and GOLPH3 bound to the membrane in assembled COPI-coated vesicles. This reveals that GOLPH3 actually makes two contacts with COPI, the first via its flexible N-terminal domain, and the second via a direct interaction with the beta-propellor domain of α-COP, occluding one of the two sites that bind to the K(X)Kxx signal in ER residents. The interaction with α-COP holds the acidic face of GOLPH3 just above the membrane surface, and mutation of membrane-proximal acidic residues results in loss of retention of Golgi residents, with different mutations affecting different residents. These findings reveal how COPI transports Golgi resident enzymes and how it switches specificity so as to recycle Golgi residents from later Golgi compartments but not from earlier ones.

### Preparation of COPI coat bound to GOLPH3 for structural studies

To characterize the interactions between the COPI vesicle coat and GOLPH3, we reconstituted vesicle formation from purified components as previously described with minor modifications to allow incorporation of GOLPH3 (*27–29*). In the minimal COPI-coated vesicle budding reaction, giant unilamellar vesicles are incubated with purified coatomer, the small GTPase Arf1, a guanine exchange factor, ARNO, and GTPγS, a nonhydrolyzable analog of GTP (fig. S1, A-D). To incorporate GOLPH3 into the assembled vesicles, purified GOLPH3 was added at a 6-7-fold excess relative to coatomer. As membrane binding of GOLPH3 is dependent on the lipid PI4P (*21*, *30*), 3 mol% PI4P was added to the lipid mixture used to prepare the GUVs. These two modifications allowed incorporation of GOLPH3 into the budded vesicles. After incubation at 37°C for 30 minutes, aliquots of the reaction mixture were applied to cryoEM grids and vitrified for characterization by cryo-electron tomography (cryoET).

### CryoET structure of GOLPH3 incorporated in the assembled COPI coat

We collected 206 tomograms of the COPI-GOLPH3 budding reaction sample and 4866 COPI vesicles and buds were manually identified. Initial particle locations for subtomogram averaging were seeded evenly over the surface of each sphere corresponding to a vesicle or bud (fig. S2A). Subtomogram averaging was performed using a low-resolution structure of the COPI triad (EMDB-2985, (*29*)) as a reference and the final aligned positions formed a regular lattice as previously described for COPI triads in COPI-coated vesicles and buds (*28*) (fig. S2B). We obtained a structure of the COPI leaf bound to GOLPH3 (one COPI heptamer in complex with two Arf1 molecules and one molecule of GOLPH3) at 7.5 Å resolution (fig. S2, C-F). At this resolution, alpha helices are clearly resolved, which allowed us to generate a model of the COPI leaf bound to GOLPH3 (Fig. 1). The positions of densities for the COPI components and Arf1 within the assembled coat are very similar to the previous model; however, there is a clear additional density that corresponds to GOLPH3 adjacent to the membrane, positioned between ζ-COP and the N-terminal β-propeller of α-COP (Fig. 1, B to D, Fig. 2A).

**Figure 1.**
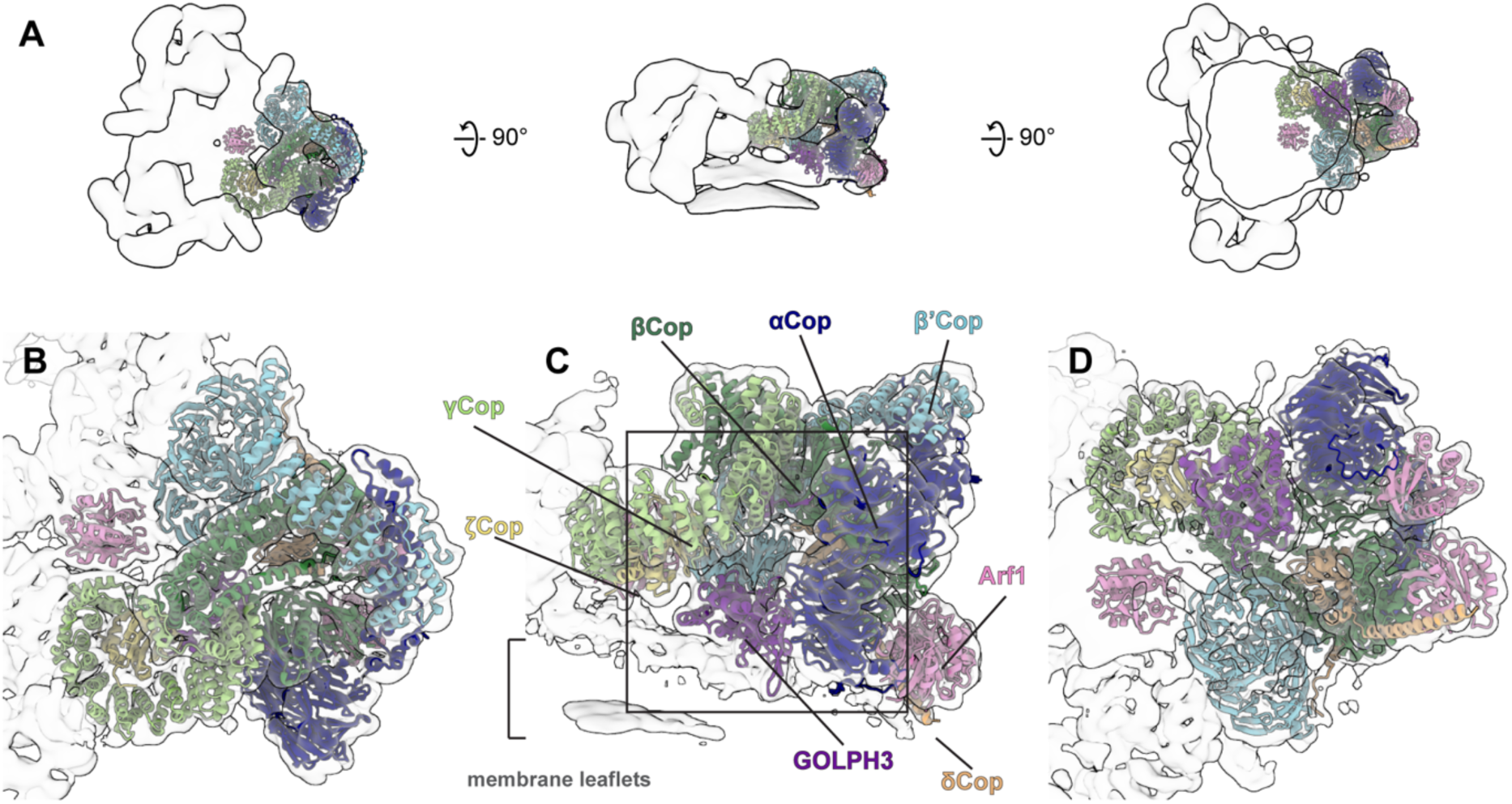
Cryo-ET structure of GOLPH3 assembled into COPI-coated vesicles. (A) Three views of the EM density map of the COPI triad, shown as an outlined semi-transparent surface to illustrate viewing directions, lowpass filtered to 20 Å. From left to right: the top view from the vesicle exterior looking down toward the membrane, the side view perpendicular to the membrane, and bottom view from below the membrane. The triad densities are fitted with ribbon models of the COPI heptameric leaf containing GOLPH3 to illustrate the location of the asymmetric unit of coatomer in the triad that constitutes the building block of the vesicle coat. (B-D) Semi-transparent isosurface of the COPI leaf EM density map fitted with ribbon models of the COPI heptameric leaf containing GOLPH3 coloured as follows: GOLPH3, purple, α-COP, dark blue; β-COP, dark green; β’-COP, light blue; δ-COP, orange; γ-COP, light green; ξ-COP, yellow, Arf1, pink. The square on panel C indicates the position of the zoomed-in view shown in Fig. 2A.

**Figure 2.**
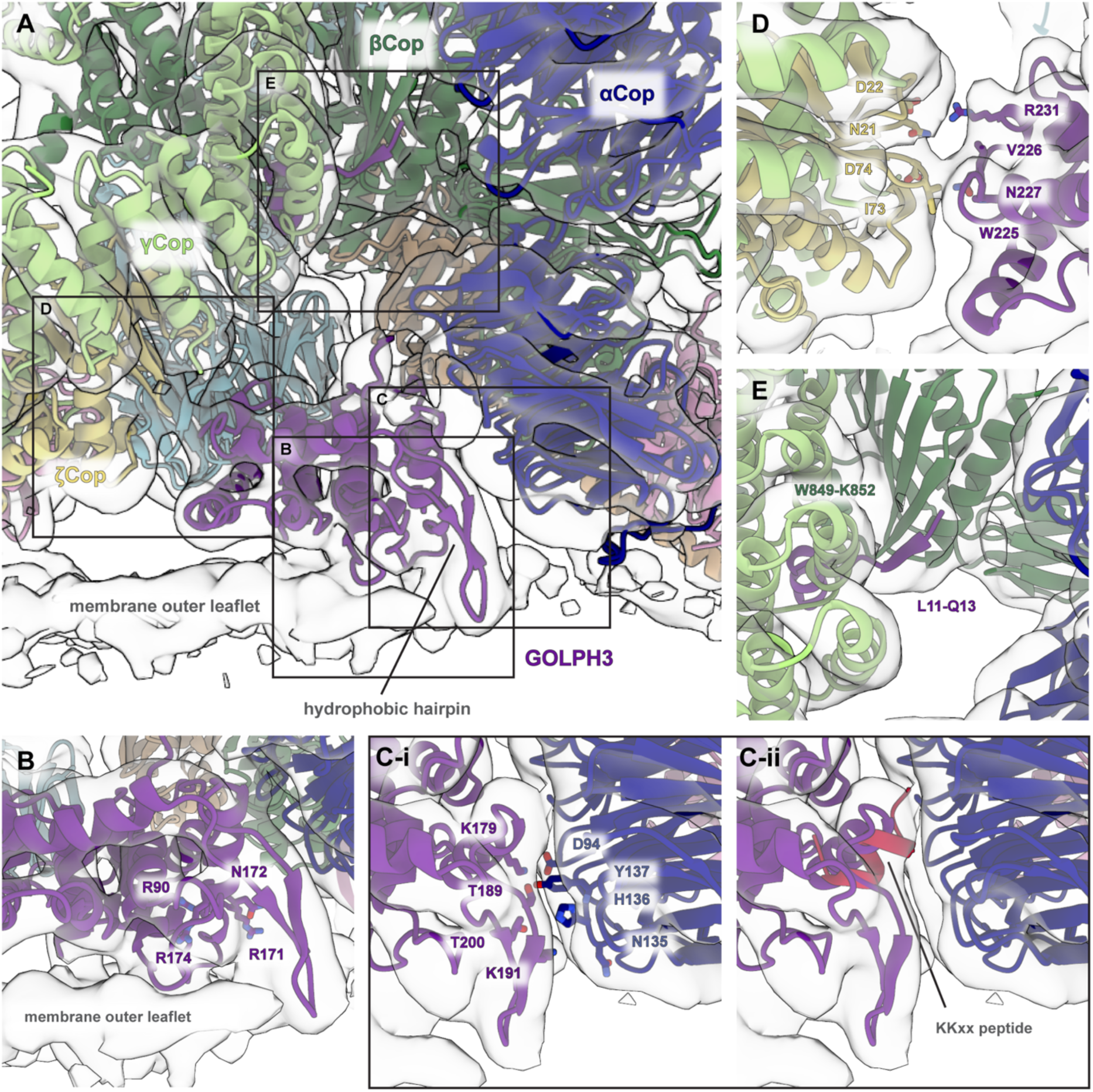
GOLPH3 is positioned proximally to the membrane, located between α-COP and ξ-COP. (A) Semi-transparent isosurface fitted with a ribbon model of the COPI-GOLPH3 leaf. The view is centered on GOLPH3 and is perpendicular to the membrane. Boxes indicate the positions of the zoomed in views shown in (B-E). (B) Zoomed in view of the PI4P binding site, with resides that mediate PI4P binding shown in sticks (R90, R171, N172, R174). (C) (i) Zoomed in view of the interface between GOLPH3 and α-COP with interacting residues indicated. (ii) GOLPH3 occupies the KKxx binding site of α-COP. The structure of the N-terminal β-propellor of α-COP bound to a KKxx peptide (E19, PDB: 4J8G) was aligned to COPI-GOLPH3 leaf model. Only the peptide (RRSFIDEKKMP, crimson) is shown, which overlaps with the position of GOLPH3. (D) Zoomed in view of the interface between GOLPH3 and ξ-COP with interacting residues indicated. (E) The N-terminus of GOLPH3 forms an extended beta sheet with β-COP. The residues involved are labelled. Residue R14 in GOLPH3 is required for interaction with coatomer and is positioned close to two negatively charged residues in γ-COP (D445 and E447). Components are coloured as follows: GOLPH3, purple, α-COP, dark blue; β-COP, dark green; β’-COP, light blue; δ-COP, orange; γ-COP, light green; ξ-COP, yellow, Arf1, pink.

Based on the EM density, we performed rigid body fitting of individual domain structures to generate a molecular model of the GOLPH3-containing coat (see Materials and Methods). In the previous model of the COPI coat, several small densities which were not filled by the known structures and homology models used to build the model were assigned using a combination of secondary structure prediction and cross-linking mass spectrometry (*27*). In our model of the COPI leaf bound to GOLPH3 the improved resolution of the model largely confirmed the identification of these densities. A density adjacent to the N-terminal beta-propeller of α-COP that was previously proposed to be constituted of a flexible loop in α-COP can now be unambiguously assigned as an additional copy of Arf1 at incomplete occupancy (fig. S3A). Although the stoichiometry of Arf1:COPI in vivo is 2:1, we believe that the third copy of Arf1 observed in the COPI leaf here may result from excess Arf1 added to the in vitro budding reaction (*9*). The resulting model allows us to interpret the interactions between GOLPH3, the COPI coat, and the membrane.

### Interactions between GOLPH3, the membrane and the COPI coat

GOLPH3 consists of a flexible N-terminal region (residues 1-58) and a folded C-terminal domain consisting of an alpha-helical core with a hydrophobic beta-hairpin that extends away from the core of the protein (*17*, *21*). In our structure of GOLPH3 incorporated into COPI-coated vesicles, the C-terminal domain of GOLPH3 is positioned in close proximity to the membrane, oriented such that the hydrophobic hairpin is inserted into the lipid bilayer (Fig. 2A). The residues that comprise the proposed PI4P binding site (R90, R171, N172, R174) are located close to the membrane interface, which would allow PI4P binding (Fig. 2B). These interactions result in a large, relatively flat, face of GOLPH3 being held close to the membrane surface.

GOLPH3 is located between the N-terminal β-propeller of α-COP and ζ-COP, with the majority of the interaction interface provided by α-COP. For the interface between GOLPH3 and α-COP, residues T189, K191, and T200 in the hydrophobic hairpin of GOLPH3 are positioned against the α-COP beta-propellor in the vicinity of residues N135, H136, and Y137 (Fig. 2C). Residues E175, K179 and E183 in GOLPH3 are positioned along one face of an alpha helix where they could interact with the loops of the α-COP beta-propellor. In particular, K179 in GOLPH3 seems well-positioned to interact with residue D94 in α-COP, while E183 may interact with R55 in α-COP (Fig. 2C). The β-propeller of α-COP is known to bind KKxx signals in the cytoplasmic tails of ER resident proteins (*12*, *13*). In our structure GOLPH3 binds over and occludes the binding pocket for KKxx motifs (Fig. 2C). The interface between GOLPH3 and ζ-COP is less extensive than that with α-COP. A loop in ζ-COP consisting of residues S72, I73, and D74 is in close proximity to a section of a loop in GOLPH3 consisting of residues D223 to N227, while residue R231 in GOLPH3 is positioned close to residues N21 and D22 in ζ-COP (Fig. 2D).

The N-terminal domain of GOLPH3 is flexible and residues 1-51 were deleted from the protein used to produce the published crystal structure (*21*). However, the N-terminal region has been proposed to be important for GOLPH3 incorporation into COPI vesicles (*20*). In each prediction of the COPI-GOLPH3 leaf structure that we performed, Alphafold3 predicted that residues 11-13 (LVQ) in GOLPH3 interact with residues W849 – K852 of β-COP to continue the beta sheet in the β-COP appendage domain of β-COP (see methods). Inspection of the EM structure revealed density corresponding to this portion of GOLPH3 at the predicted position (Fig. 2E). Consistent with this, R14 in human GOLPH3, adjacent to this motif and in the vicinity of D445 and E447 in γ-COP, has been shown to be required for interaction with coatomer (*20*), and the leucine and arginine of the LVQR sequence are very well conserved in GOLPH3 orthologues from diverse species including yeasts (fig. S5A).

The interactions that form between GOLPH3 and α-COP result in subtle confirmational changes in GOLPH3 and in the assembled COPI coat. The GOLPH3 hydrophobic hairpin is positioned slightly closer to the alpha helical core of the protein in the EM-density relative to the crystal structure of GOLPH3 (fig. S3). The α-COP N-terminal beta-propeller domains are better resolved relative to the rest of the complex when compared to the previous structure of the coat without GOLPH3, suggesting that the position of the α-COP beta-propellors within the coat is stabilized by GOLPH3.

### Interaction between GOLPH3 and α-COP is required for function

In cell lines lacking GOLPH3 and its less abundant paralogue GOLPH3L, many Golgi resident proteins are mislocalised with a concomitant reduction in abundance as they are either trafficked to the lysosome for degradation or are clipped by proteases such as SPPL3 and subsequently secreted (*19*, *24*, *31*) (Fig 3A). Thus, to test the functional significance of interactions between GOLPH3 and coatomer, we used immunoblotting and immunofluorescence microscopy to monitor the ability of mutated GOLPH3 proteins to rescue the levels and localisation of Golgi residents in the GOLPH3/3L deletion cell line (Fig 3A and fig. S4). Two important functional regions of GOLPH3 are the PI4P binding site and the hydrophobic beta-hairpin that has been proposed to insert into the membrane, and mediate dimerization of soluble GOLPH3 (*21*, *32*). These previous studies have shown that mutation of either of these regions results in a loss of GOLPH3’s Golgi localisation and its resident protein retention activity, and we could recapitulate this using our in vivo assay (fig. S5, B and C). GOLPH3 with mutations K179E and E183K in the interface between GOLPH3 and the α-COP β-propellor domain was stable when expressed in the GOLPH3/3L cell line but was unable to rescue the mislocalisation of Golgi residents known to depend on GOLPH3 for their localisation (Fig. 3, B to E). Interestingly, although inactive for retention, the mutant GOLPH3 protein itself was still localised to the Golgi (Fig. 3C). In contrast, GOLPH3 with the mutations N227A and R231A in the interface with ζ-COP could rescue both function and Golgi localisation of GOLPH3 (fig. S5, D and E). Thus, the key functional interaction that the C-terminal folded domain of GOLPH3 makes with the COPI coat appears to be that with α-COP.

**Figure 3.**
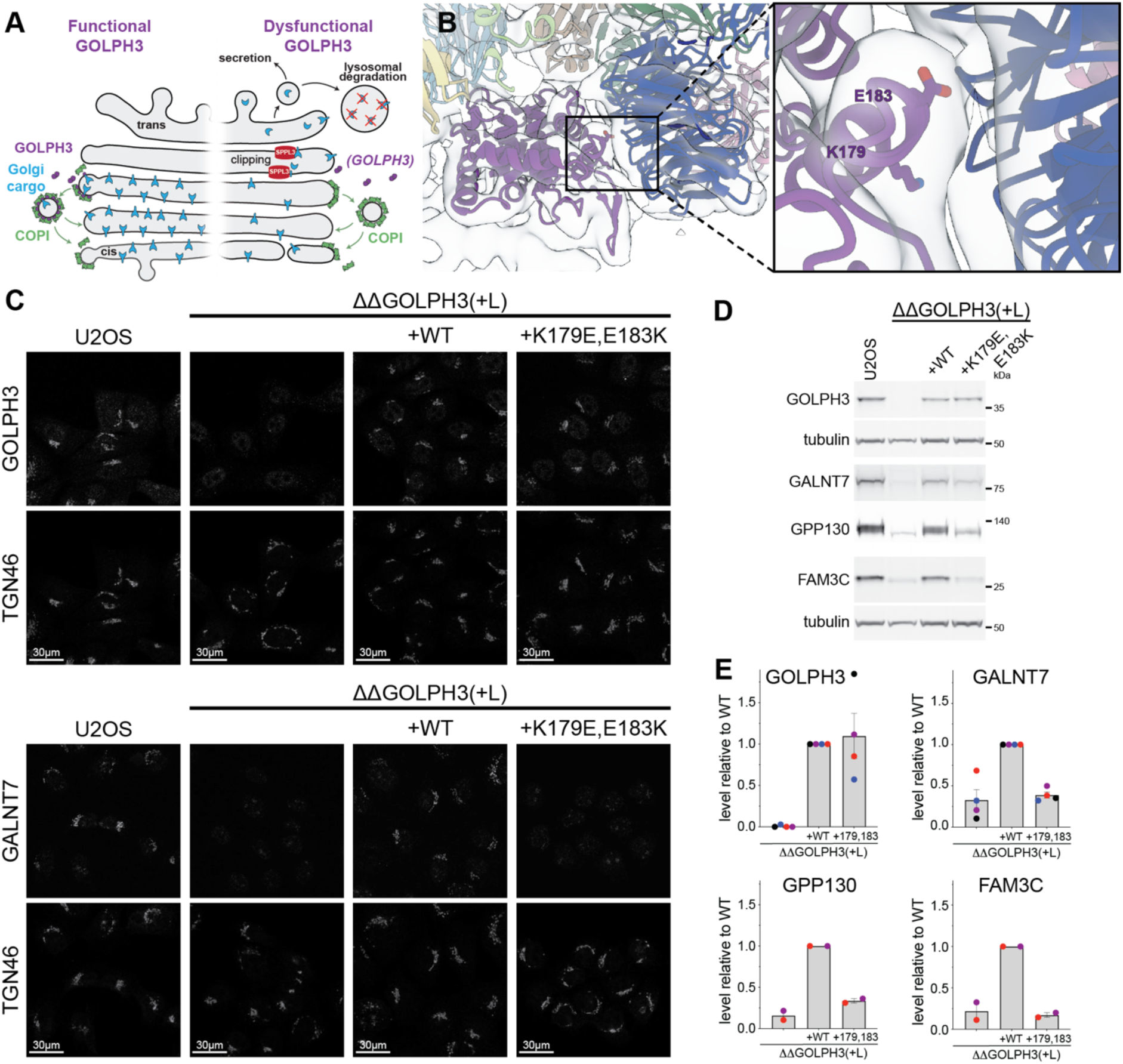
Interactions at the interface between GOLPH3 and α-COP are required for proper function of GOLPH3 in cargo localisation. (A) Loss of GOLPH3 activity destabilises a subset of Golgi residents as they are no longer retained and instead are either secreted having been clipped in later compartments by Golgi proteases such as SPPL3 or are trafficked to the lysosome for degradation (*19*, *24*, *31*). (B) The interface between GOLPH3 and α-COP with interface residues K179 and E183 shown as sticks in the zoomed panel (right). (C) Confocal micrographs of U2OS cells labelled for the Golgi marker TGN46 and either GOLPH3 or the Golgi resident enzyme GALNT7. The cells are either wild type or lacking both GOLPH3 and GOLPH3L with the latter mutant transfected with either wild-type GOLPH3 or a version with K179E and E183K mutations. (D) Immunoblot of whole-cell lysates of the cells shown in (C) and labelled for the indicated Golgi resident proteins or tubulin as a loading control. FAM3C is type II Golgi protein of unknown function (*48*, *49*). (E) Quantification of the levels of the indicated proteins from the immunoblot shown in (D) and independent replicates of this blot. Mutation of residues K179 and E183 does not perturb the levels or Golgi localisation of GOLPH3, but prevents it being able to rescue retention of the Golgi residents.

Taken together, these results suggest that the initial recruitment of GOLPH3 to Golgi membranes is via a combination of binding of the N-terminus to COPI, binding to PI4P, and insertion of the hydrophobic hairpin. The interaction with the α-COP beta-propellor is then required for GOLPH3 to adopt a functional conformation within the coat.

### Interactions with Golgi resident proteins are mediated by distinct regions of the GOLPH3 membrane-adjacent surface

GOLPH3 has a highly electronegative face that has been previously proposed to interact with cytoplasmic tails of glycosylation enzymes which are generally short and highly basic (*19*). In our structure, this surface faces the membrane with the hydrophobic hairpin and the PI4P binding contacting the membrane whilst the rest rises slightly above the membrane surface to create a wedge-shaped chamber with the electronegative surface as its roof (Fig. 4A). To explore the role of this electronegative surface we tested the effect of mutating clusters of acidic residues to alanine on GOLPH3 activity (Fig. 4A and B). We first assessed the stability and localization of GOLPH3 containing each cluster of mutations. Although one pair of overlapping clusters destabilised GOLPH3 (B and B’), GOLPH3 containing any of four other mutant clusters (A, A’, C and D) was stable and localised to the Golgi when expressed in the GOLPH3/3L deletion cell line indicating that it is correctly folded (Fig. 4C and D, and fig. S6A).

**Figure 4.**
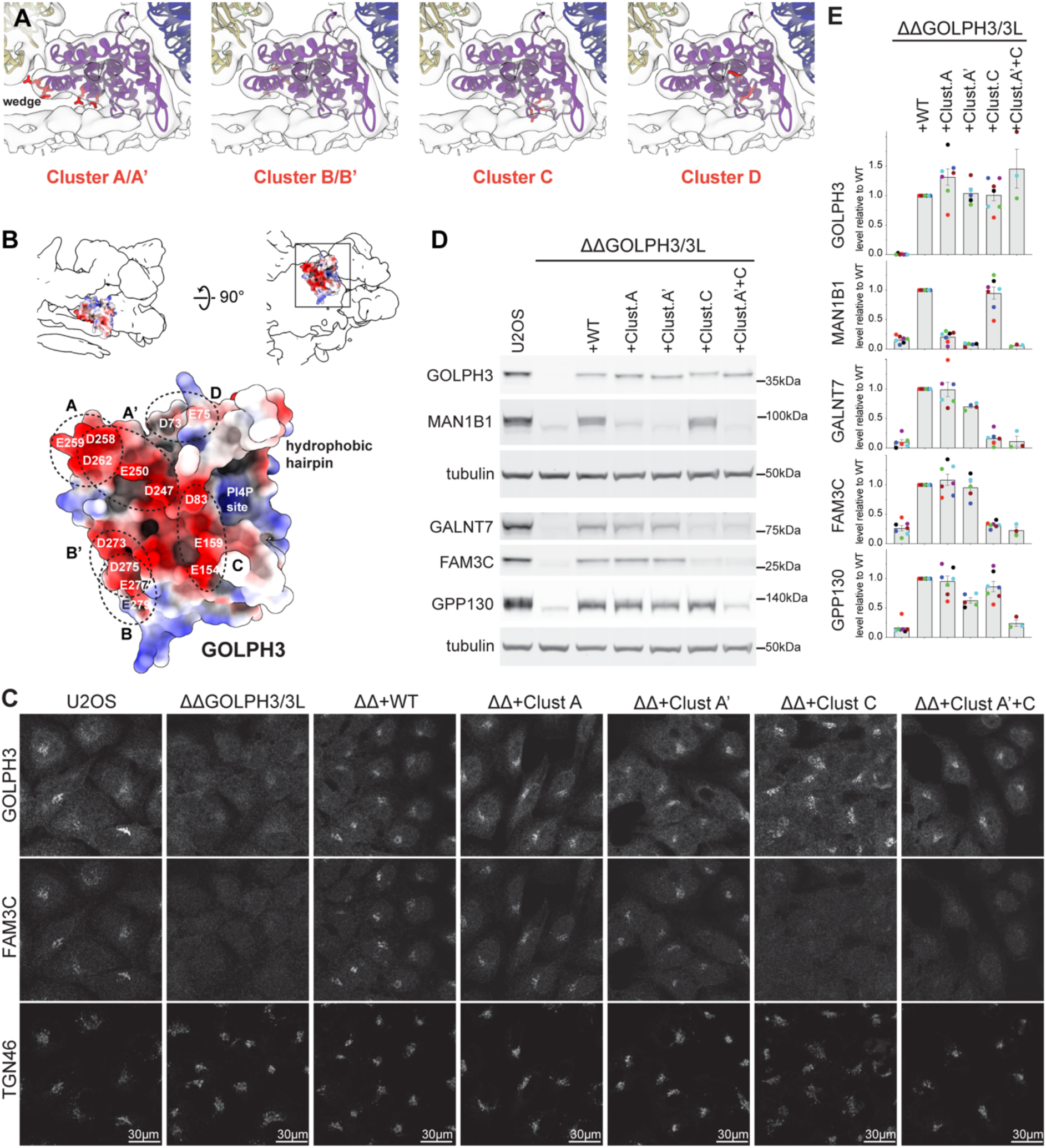
Interactions between GOLPH3 and cargo are mediated by clusters of positively charged residues. (A) A view of the semi-transparent COPI-GOLPH3 leaf density perpendicular to the membrane with ribbon models fitted. Clusters of mutated residues are labelled in red to show their position relative to the membrane. (B) The model of GOLPH3 is shown as a surface coloured by electrostatic charge and fitted into the lowpass filtered (15 Å) COPI-GOLPH3 leaf density, delineated as a silhouette. A view perpendicular to the membrane is on the left and a 90° rotation viewed from the membrane surface is on the right. Below, the membrane-facing side of GOLPH3, shown as a surface coloured by electrostatic charge, with the clusters of mutated residues indicated. (C) Confocal micrographs of U2OS cells labelled for the Golgi marker TGN46 and GOLPH3 and the GOLPH3-dependent Golgi resident protein GPP130. The cells are either wild type or lacking both GOLPH3 and GOLPH3L (ΔΔGOLPH3/3L (ΔΔ)) with rescue by either wild type GOLPH3 (WT) or mutants in which the clusters of acidic residues shown in (A and B) have been mutated to alanine. (D) Immunoblots of whole cell lysates from U2OS cells as in (C) probed for the indicated Golgi residents, or for tubulin as a loading control. (E) Quantification of the levels of the indicated Golgi proteins in the immunoblot shown in (D) and replicates of this blot. For each experiment, the levels of each protein were normalised to the level expressed when wild-type GOLPH3 was transfected back into the GOLPH3 and GOLPH3L double deletion cell line (ΔΔGOLPH3/3L+WT). The different clusters of mutations affect the retention of different sets of residents.

We next examined the stability and localization of a panel of different GOLPH3 cargoes (GPP130, GALNT7, MAN1B1, and FAM3C) in each mutant GOLPH3 background. GOLPH3 mutated in overlapping clusters A and A’ maintained levels of Golgi residents GALNT7 and FAM3C but had reduced levels of MAN1B1 relative to the wild type (Fig. 4D and E). In contrast, mutations in cluster C gave reduced levels of GALNT7 or FAM3C but maintained the levels of MAN1B1 relative to wild type. GOLPH3 with mutations in Cluster D rescued the levels of all tested proteins (fig. S6B). GPP130 retention was not affected by any of the individual mutant clusters, but GOLPH3 with mutations in both clusters A’ and C no longer rescued the location or levels of GPP130 despite being stable and Golgi-localized (Fig. 4C-E; fig S5C). Quantification of replicates showed that these differences were highly reproducible (Fig. 4E). This striking finding indicates that not only does the acidic surface of GOLPH3 act directly in retention of Golgi enzymes but that different parts of the GOLPH3 acidic surface are involved in the recognition of different clients.

## Conclusions

The organisation of trafficking within the Golgi has long been controversial (*8*, *33*). Debate initially focused on whether secretory cargo is transported through the stack in anterograde vesicles moving between stable cisternae or whether the cisternae form at the cis side and maturing to the trans-side whilst retrograde vesicles recycle Golgi residents. A key step toward understanding the molecular basis of transport came when the COPI coat was identified through an in vitro reconstitution of vesicle-mediated transport between Golgi stacks (*34*, *35*). This assay was interpreted as reconstituting anterograde vesicle transport, but this interpretation was called into question by the subsequent discovery that COPI is responsible for retrograde traffic of escaped ER-residents from the cis Golgi to the ER (*36*). As a result, transport within the stack has continued to be debated with various models proposed for how COPI might mediate either retrograde or anterograde traffic (*7*, *37–40*). The results presented here shed light on two long-standing questions in this debate. First, they imply a clear molecular model for how the COPI coat recruits resident Golgi enzymes for retrograde transport within the Golgi. Second, the results allow us to propose how COPI-coated vesicles can mediate two distinct transport routes, switching from their well-established role in retrieval from the cis Golgi of escaped ER residents to take on a second role of recycling Golgi residents from later cisternae.

GOLPH3 is known to form homodimers in solution through its hydrophobic hairpin but when assembled in the COPI coat GOLPH3 is monomeric with the hairpin buried in the bilayer. Thus, dimerization seems to allow GOLPH3 to be soluble in the cytoplasm until it partitions into membranes, as was predicted by earlier membrane binding studies (*32*). Interactions between the GOLPH3 N-terminus and β-COP, between the GOLPH3 C-terminal domain and the α-COP propeller, and binding of GOLPH3 to PI4P, stabilise the interaction within the assembling coat and position GOLPH3 close to the membrane. As described above, GOLPH3 is tilted to form a wedge-shaped chamber above the bilayer that is closed at one end by contacts with the bilayer through PI4P and the hydrophobic hairpin. The acidic residues line the roof of this chamber with those in cluster C being close to the bilayer at the closed end of the chamber (Fig. 4A). These residues are required for retention of GALNT7 and FAM3C, both of which have short basic tails. In contrast, the acidic residues in clusters A and A’ are near the entrance to the chamber and further from the bilayer, and these residues are required for retention of MAN1B1 which has an unusually long cytoplasmic tail of 48 residues. Thus, we speculate that GOLPH3 selects proteins with short basic tails as only these can reach cluster C at the back of the chamber, but it can also bind specific clients with longer tails via sequence-specific interactions with residues near the entrance to the chamber. A diversity of binding mechanisms, combined with the use by different clients of different parts of the GOLPH3 surface would provide an explanation for why a consensus sequence for tail-mediated Golgi retention has proven hard to define.

COPI-coated vesicles initially form on the earliest post-ER structures – the ER-Golgi intermediate compartment (ERGIC) - and their formation continues throughout the Golgi stack (*41*, *42*). Our results allow us to propose how the cargo specificity of COPI-coated vesicles adapts from retrieval of escaped ER residents at the ERGIC and cis Golgi to also allow recycling of Golgi residents in the later cisternae (Fig. 5). PI4P levels at the ERGIC and cis Golgi are low because PI4P is made later in the stack, and because the PI4P phosphatase Sac1 recycles between the ER and the cis Golgi (*43–45*). Consistent with this, high-resolution imaging and organelle proteomics has shown GOLPH3 is present on the later Golgi cisternae rather than the ERGIC and cis Golgi (*46*, *47*). Thus, GOLPH3 incorporation into COPI-coated buds is not favoured at the ERGIC and cis Golgi and so only ER residents are recruited by interacting with the terminal beta propellors of α-and β’-COP via KKxx and other signals. In addition, the presence of larger numbers of KKxx-containing proteins binding the α-COP beta propellor would occlude the main site of interaction between GOLPH3 and coatomer, further preventing recruitment of GOLPH3 to the cis Golgi. As the Golgi matures, the levels of PI4P will increase as the kinase is recruited and the Sac1 phosphatase is recycled back to the ER. This increase in the levels of PI4P will promote the incorporation of GOLPH3 into COPI-coated buds and hence the recruitment of Golgi residents. GOLPH3 incorporation in COPI-coated vesicles will block the KKxx cargo binding site at the α-COP beta propellor, but the binding site at the β’-COP propellor would remain free to mediate transport of any remaining ER-destined cargo. This scenario would allow spatial separation of the GOLPH3-COPI mediated transport of Golgi resident enzymes in the later Golgi and the COPI mediated transport of ER resident proteins away from the cis-Golgi, while including a fail-safe mechanism for retrograde transport of ER residents that are mislocalised to the later Golgi cisternae. Further mechanisms may exist to allow COPI to recruit additional clients such as membrane traffic machinery or enzymes that are located to specific regions of the stack, but our work makes clear that COPI is able to perform at least two distinct roles in recycling thus providing a mechanistic basis for the Golgi maturation model.

**Figure 5.**
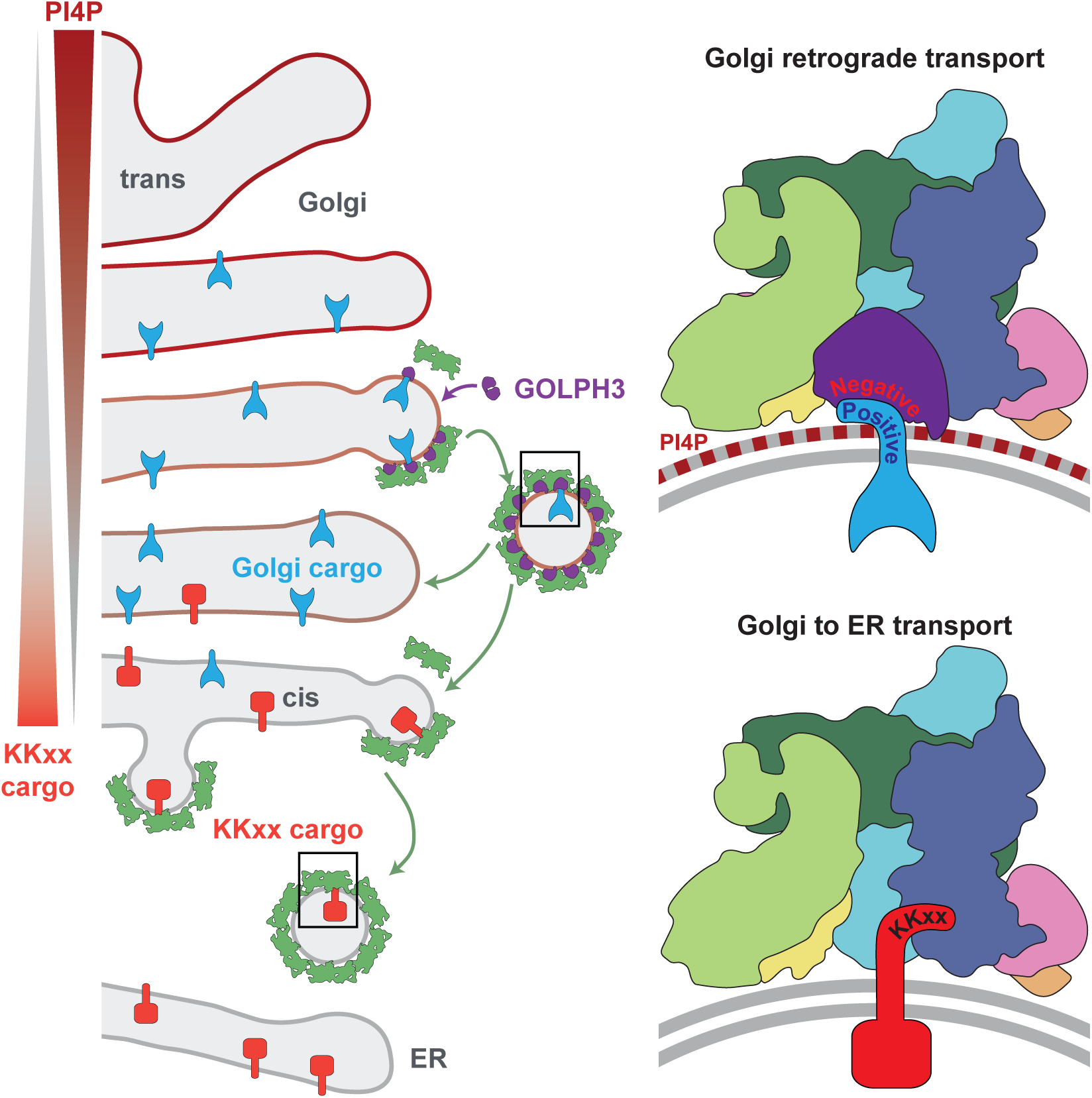
GOLPH3-mediated selection of COPI vesicle cargo during Golgi maturation. A model for how COPI can switch its cargo specificity during Golgi maturation. In the cis Golgi, COPI is recruited by Arf1 and ER proteins with dilysine motifs (KKxx) bind to the beta propellor of α-COP for recycling to the ER. GOLPH3 does not bind as levels of PI4P are low (due the trans Golgi location of PI4KB and the action of the phosphatase Sac1 in the cis Golgi), and because KKxx cargoes occupy the GOLPH3 binding site in the beta propellor of α-COP. As the Golgi matures, levels of PI4P increase and Sac1 is removed, while the concentration of KKxx cargo is reduced. This directs integration of GOLPH3 into the COPI coat where it binds the beta propellor of α-COP such that its negatively charged surface is held close to the membrane to recruit Golgi residents by their short positively charged tails. Any remaining proteins with a KKxx motif would still be able to bind to the beta propellor of β’-COP. Thus, COPI vesicles recycle Golgi residents to earlier Golgi cisternae where they can function as the cisternae matures until the next round of recycling.

## Acknowledgments

We thank Patrick Aderhold, Iva Ganeva, Simone Pienz, Lawrence Welch and Felix Wieland for providing reagents, Guiseppe Cannone for assistance with data collection; Florian Beck and Inga Wolf for assistance with computing infrastructure; and Manu Hegde for comments on the manuscript. This study made use of electron microscopes at the MRC-LMB EM Facility and the high-performance computing resources at the Max Planck Institute of Biochemistry.

## Author contributions

Conceptualization JAGB, SM, DJO

Methodology: JAGB, KAC, GT, RJT

Investigation: KAC, JGGK, DJO, RJT, NZ

Resources: JGGK, NZ

Visualization: RJT, NZ

Funding acquisition: JAGB, SM, DJO

Supervision: JAGB, SM

Writing – original draft: JAGB, SM, RJT

Writing – review and editing: JAGB, SM, DJO, RJT, NZ

## Data and materials availability

The structure determined by electron microscopy is deposited in the Electron Microscopy Data Bank under accession code EMD-XXXXX. The corresponding molecular model is deposited in the Protein Data Bank under accession code XXXX.

## SUPPLEMENTARY MATERIALS

Materials and Methods

Figs. S1 to S7

Tables S1 and S2

## Materials and Methods

### Protein purification for reconstitution of COPI budding

Recombinant coatomer was prepared in the lab of Felix Wieland and was purified as previously described (*50*). Briefly, *M. musculus* coatomer isoforms γ1ξ1, γ1ξ2, and γ2ξ1 were each recombinantly expressed in SF9 insect cells using a baculoviral expression system. The recombinant coatomer was affinity purified using a One-Strep-Tag at the C-terminus of α-COP followed by purification by size exclusion using a Superose 6 10/300 gl column (Cytiva) in buffer containing 25 mM HEPES pH 7.4, 200 mM KCl, 10% glycerol, and 1 mM DTT. After separate purification of each isoform, the final coatomer sample used in budding reactions was prepared by mixing the isoforms at a ratio of 2:1:1 for γ1ξ1:γ1ξ2:γ2ξ1. Myristoylated human Arf1 and human ARNO was prepared as described previously (*51*, *52*).

Full-length human GOLPH3 was expressed recombinantly in BL21(DE3) pLysS *E. coli* grown in 2xTY media, with expression induced with 0.2 mM IPTG at 22 °C for 16 hours. Cells were resuspended in buffer A (300 mM NaCl, 20 mM Tris, pH 7.4, 1 mM DTT) supplemented with AEBSF hydrochloride, MnCl_2_, and DNAseI. The supernatant was batch bound to glutathione Sepharose resin (Cytiva) in buffer A, the resin was washed with 400 mL of buffer A, and the protein was eluted by thrombin cleavage of the N-terminal GST tag. The resultant flow through was purified by Ni-NTA purification using a N-terminal His_6_ tag. Eluted fractions were concentrated for gel filtration Superdex 200 HiLoad 26/60 column (Cytiva) into buffer containing 10 mM Tris pH 7.4, 350 mM NaCl, and 1 mM DTT. Pooled and concentrated fractions were supplemented with AEBSF and 10 mM DTT before freezing.

### Preparation of liposomes

All lipids were purchased from Avanti Polar lipids. Liposomes were prepared using a lipid mixture consisting of 34 mol % DOPC, 30 mol % DOPE, 15 mol % DOPS, 15 mol % DOPA, 3 mol % PI4P, 5 mol % PIP_2_, and 1 mol % of the fluorescent dye DiI (Invitrogen). This lipid mixture contains an increased proportion of acidic phospholipids, as this has been observed to promote formation of COPI-coated vesicles and contains 3 mol % PI4P to allow recruitment of GOLPH3 (*21*, *30*, *53*).

The lipid mixture was dried under vacuum overnight and was resuspended at a concentration of 2.3 mg/ml in 50 mM HEPES pH 7.4, 100 mM KOAc, and 1 mM DTT with 1 mM MgCl_2_ added after rehydration of the lipid film. Liposomes were extruded through a 400 nm polycarbonate membrane (Avanti Polar Lipids) to generate unilamellar liposomes with a uniform size distribution.

### In vitro budding reaction and cryoET sample preparation

COPI-coated vesicles with GOLPH3 incorporated in the coat were prepared by an *in vitro* budding reaction, similar to that previously described (*27–29*), with minor modifications. Liposomes (0.23 mg/ml) were incubated with GTPγS (0.5 mM), COPI (1.36 μM), Arf1 (20 μM), ARNO (4 μM), and GOLPH3 (9 μM) in a total volume of 20 μL in buffer consisting of 50 mM HEPES pH 7.2, 125 mM KOAc or acetate, and 2.5 MgCl_2_. After incubation for 30 minutes at 37°C, samples were prepared for cryoET by plunge freezing in liquid ethane using a manual plunger. Reaction mixture (3 μL) was applied to glow-discharged CF2/2 Cu300 grids (C-Flat) and the grids were manually blotted from the back for 5 s before freezing.

### Data acquisition

Tilt series were collected using a 300 kV Titan Krios microscope (ThermoFisher Scientific) equipped with a Quantum K3 direct electron detector (Gatan) and a Quantum energy filter (Gatan) with a slit width of 20 eV. The tilt series were collected in counting mode using SerialEM-3.8.0 software, at a pixel size of 1.71 Å with a dose symmetric scheme with an angular range of -60° to +60° in 3° increments (*54*, *55*). The total dose was 125 e^-^/Å^2^, evenly distributed across tilts, with each tilt image acquired as a 10-frame movie. The applied defocus ranged from -1.5 to -3.5 μm.

### Tomogram reconstruction

Raw movies were motion-corrected and gain-corrected in IMOD-4.11.19 and aligned stacks were generated using the alignframes function in IMOD (*56*). A dose-filtered stack with each image in the tilt series low-pass filtered based on the accumulated electron dose during acquisition (*57*) was also generated using the alignframes function in IMOD. The contrast transfer function (CTF) was estimated using the ctfplotter function in IMOD on the non-dose-filtered stack (*58*). Tilt series stacks were aligned using fiducial-based alignment in IMOD and bin8 tomograms without CTF correction were reconstructed in IMOD by weighted back projection. For initial particle picking and better visualization, tomograms were low-pass filtered to 50 Å using EMAN2.99.47 (*59*). CTF-corrected tomograms were generated using novaCTF (*60*).

### Subtomogram averaging

A total of 4866 COPI vesicles and buds were manually picked from 206 tomograms in UCSF Chimera (*61*), with the tomograms binned 8 times to a pixel size of 13.6Å. The center of each coated vesicle or bud was picked and radii were assigned using a custom-written plugin (https://www.biochem.mpg.de/7940000/Pick-Particle (*62*)). Initial coordinates for subsequent particle search were seeded with an even spacing of 10.9 nm (8 pixels in tomograms binned 8 times) perpendicular to the sphere surface, and with random in-plane rotation, yielding a total of 193104 particles.

Bin8 subtomograms (48 pixel box size) were extracted and iteratively aligned against the published 3-fold COPI trimer density map low pass filtered to ∼40 Å (EMDB 2985, (*29*)) using the subTOM package (v1.1.5) (https://subtom.readthedocs.io/ or https://github.com/DustinMorado/subTOM). After alignment, duplicates were removed and incorrect picks were cleaned based on the cross-correlation score and manually, resulting in a final set of 36144 particles. These particles were subsequently split into two half sets for further refinement.

Aligned tilt series and particle coordinates were transferred to Warp (v1.0.9) (*63*), where CTF estimation was performed, and subtomograms and corresponding per-particle CTF models extracted at bin2 (3.40 Å pixels size). Subsequent refinement of COPI trimer was conducted in RELION (v3.0) yielding a 12.4 Å structure (*64*).

Particles with refined coordinates and Euler angles were used to refine the tilt series alignment in M (v1.0.9) (*65*). New subtomograms were extracted at bin1.5 (2.55 Å pixel size) and particle positions were refined in RELION yielding a 9.4 Å structure. These particles were transferred to M for another round of refinement of the tilt series alignment.

After expanding C3 symmetry, new subtomograms were extracted at bin1 (1.70 Å pixel size) centered on each asymmetric COPI leaf (yielding total 108432 particles), and particle positions were refined in RELION, yielding an 8 Å structure. These particles were then transferred to M for further refinement of tilt series alignment and CTF models, resulting in the final 7.5 Å structure (table S1).

### Model building

To facilitate building a molecular model for the COPI leaf with GOLPH3, we used Alphafold 3 (AF3, (*66*)) to generate structures of a protein complex containing the following components: full-length *H. sapiens* GOLPH3 (Q9H4A6), two copies of full-length *H. sapiens* Arf1 (P84077), *M. musculus* α-COP (Q8CIE6) residues 1-900, full-length *M. musculus* β-COP (QW9JIF7), full-length *M. musculus* β’-COP (O55029), *M. musculus* δ-COP (Q5XJY5) residues 1 – 273, *M. musculus* γ1-COP (Q9QZE5) residues 1 – 609, and full-length *M. musculus* ζ-COP (P61924) (Abramson et al. 2024). This set of components represents all protein domains that are resolved in the COPI leaf. 25 Alphafold 3 runs were performed, resulting in a total of 125 models for the complex. Of these, 87 (70%) have a compressed conformation that is a poor match for the density resolved by cryoET (fig. S7A). A further 38 (30%), have a structure which is sufficiently similar to the density resolved by cryoET to be used as a basis for model building, recapitulating the observed domain interactions (fig. S7B). Notably, in 100% of the predictions, Alphafold3 predicted that residues 11-13 (LVQ) in GOLPH3 interact with residues W849 – K852 of β-COP. From this starting model of the COPI leaf, each individual protein was extracted and fit independently as a rigid body into the experimental density. For proteins containing multiple domains, each domain was separately fitted into the density as a rigid body as follows: the beta-propellor domains of α-COP (residues 1-591), the alpha solenoid domain and part of the C-terminal flexible region of α-COP (residues 592 – 824), the beta-propellor domains of β’COP (residues 1-586), the alpha solenoid and flexible C-terminal region of β’-COP (residues 587-838), the N-terminal portion of the trunk domain of β-COP (residues 1-311; with residues 660-710 which interact closely with this region of the trunk domain), the C-terminal portion of the trunk domain (residues 311 – 659), the β-COP appendage domain (residues 711 – 953; with residues 32-54 of GOLPH3 which interact closely with the β-COP appendage domain), the N-terminal portion of the trunk domain of γ1-COP (residues 1-299), the central portion of the trunk domain of γ1-COP (residues 300-488), and the C-terminal portion of the trunk domain of γ1-COP (residues 449 – 582). All rigid body fitting was performed in ChimeraX (Goddard et al. 2018, Pettersen et al. 2021, Meng et al. 2023). The regions of the coatomer complex which are not included in the model are: α-COP residues 825-1224, β’-COP residues 839-905, β-COP residues 500-535, δ-COP residues 178-223 and residues 247 - 511, γ1-COP residues 1-16 and residues 583-874, and all of ε-COP. These regions are either flexible loops/linkers or constitute the linkages between COPI triads and are not resolved in the COPI heptamer leaf structure.

### Cell culture and generation of cell lines

U2OS cells were cultured in Dulbecco’s Modified Eagle Medium (DMEM) supplemented with 10% foetal bovine serum (FBS) at 37℃ and 5% CO_2_, with regular testing to ensure that they were mycoplasma free (Lonza MycoAlert). A PiggyBac compatible expression vector with the wild-type GOLPH3 coding sequence was used for mutagenesis (*19*). COPI-binding mutants were made by sequential mutagenesis where mutations were encoded in the primers used in PCR for Gibson assembly.

Clusters of acidic residues were mutated using synthetic DNA fragments containing all desired mutations (Twist Bioscience), with the fragments inserted into the PiggyBac expression vector by Gibson assembly. A U2OS GOLPH3/GOLPH3L knockout cell line was used to stably integrate GOLPH3 variants into the genome for stable expression under the cumate-inducible promotor (*19*). Cells were seeded in a 6-well plate or a 25 cm^2^ flask. When cell reached ∼80% confluency, they were transfected with PiggyBac-compatible expression vector and the PiggyBac transposase added at an amount equalling to 500 ng of expression vector and 200 ng of transposase per well of a 6-well plate. 72 hours after transfection, cells were selected using 5 µg/ml puromycin until the cell death process stopped. After that, cells were cultured in cell culture medium containing 1 µg/ml puromycin. GOLPH3 variants expression was induced for 3 days prior to experiments by addition of 30 µg/ml of cumate to the culture medium.

### Mammalian cell lysis and immunoblotting

U2OS cells were grown in the 10cm dishes for western blot analysis, and washed once with ice-cold PBS and then scraped off the plate within PBS with 1mM EDTA and spun down at 800 x g. Cell pellet was resuspended in 4x times the volume of ice-cold lysis buffer (0.5% Triton X-100, 50 mM Tris-HCl, pH=7.4, 150 mM NaCl, 1 mM EDTA, cOmplete protease inhibitors, AEBSF). Cell lysis was performed on ice for 10 minutes before spinning down the cell debris at 16,100 x g for 10 minutes.

Bradford assay was performed on the cell lysates of U2OS cell lines to ensure equal loading of various samples onto the gel. Samples were mixed with Laemmli buffer containing 5% BME and were incubated at 90℃ for 5 minutes before loading onto gel. 4-12% gradient bis-tris gels were run in MOPS buffer at 100-175 V until the dye front reached the bottom of the gel. Gels were subjected to a western blot in which protein was transferred onto a 0.45 µm nitrocellulose membrane using a Mini Trans-Blot Cell (Bio-Rad), in transfer buffer in the presence of an ice block for 1h at a constant current of 300 mA. Blots were blocked in 5% w/v nonfat dry milk in PBST (0.5% Tween-20 in PBS) for 1 h at room temperature and incubated with the primary antibody diluted in 5% milk in PBST overnight at 4°C (table S2). Blots were washed 3 times for 5 min in PBST and incubated with the fluorescently labelled secondary antibody diluted in 5% milk in PBST for 1 h at room temperature with agitation. Blots were washed 3 times for 5 min in PBST and imaged using a ChemiDoc (Bio-Rad).

### Immunofluorescence microscopy

Cells were seeded on poly-lysine treated coverslips and three days after induction of protein expression with cumate or an appropriate time after transfection, coverslips were washed in PBS and fixed in 4% PFA for 20 minutes followed by permeabilisation with 0.5% Triton X-100 in PBS for 10 minutes. After a PBS wash, cells were subjected to a blocking buffer (0.5% Tween, 20% FBS in PBS) for 1 hour. Primary antibodies were diluted in blocking buffer at 1:200 concentration from their commercial stock solution and this solution was applied onto cells for 1 hour after the blocking step was finished. After incubation with primary antibodies (table S2), coverslips were washed with PBS and treated for 1 hour with a blocking buffer solution with fluorescent secondary antibodies at 1:300 dilution. After washing off the solution with PBS, coverslips were mounted on slides using Vectashield and sealed with nail vanish. Coverslips were imaged at room temperature using the 63x 1.4 NA oil-immersion objective on a Zeiss LSM780 confocal microscope. Brightness of images was adjusted without altering gamma or overexposing them using Fiji.

---

**Supplementary Figure 1.**
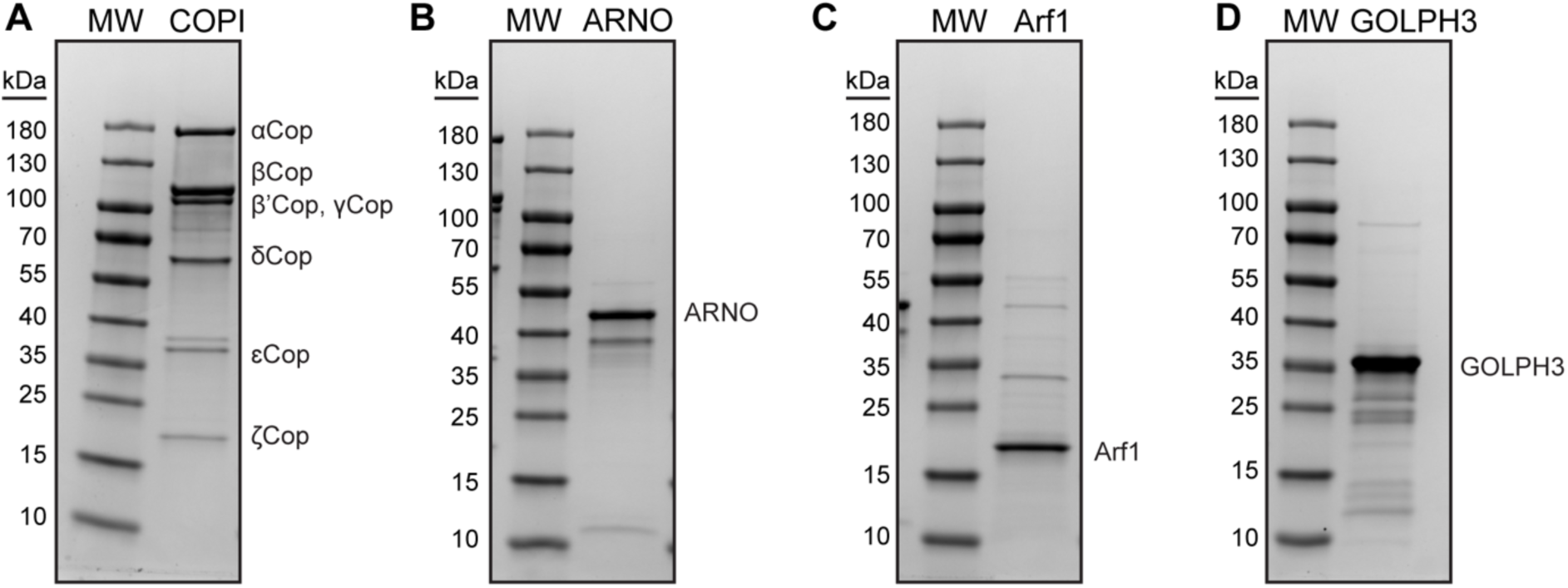
Protein preparation for in vitro reconstitution of COPI-GOLPH3 coated vesicles from purified components. (A-D) Coomassie gels for the protein components of the *in vitro* budding reaction: COPI (A), ARNO (B), Arf1 (C), and GOLPH3 (D).

**Supplementary Figure 2.**
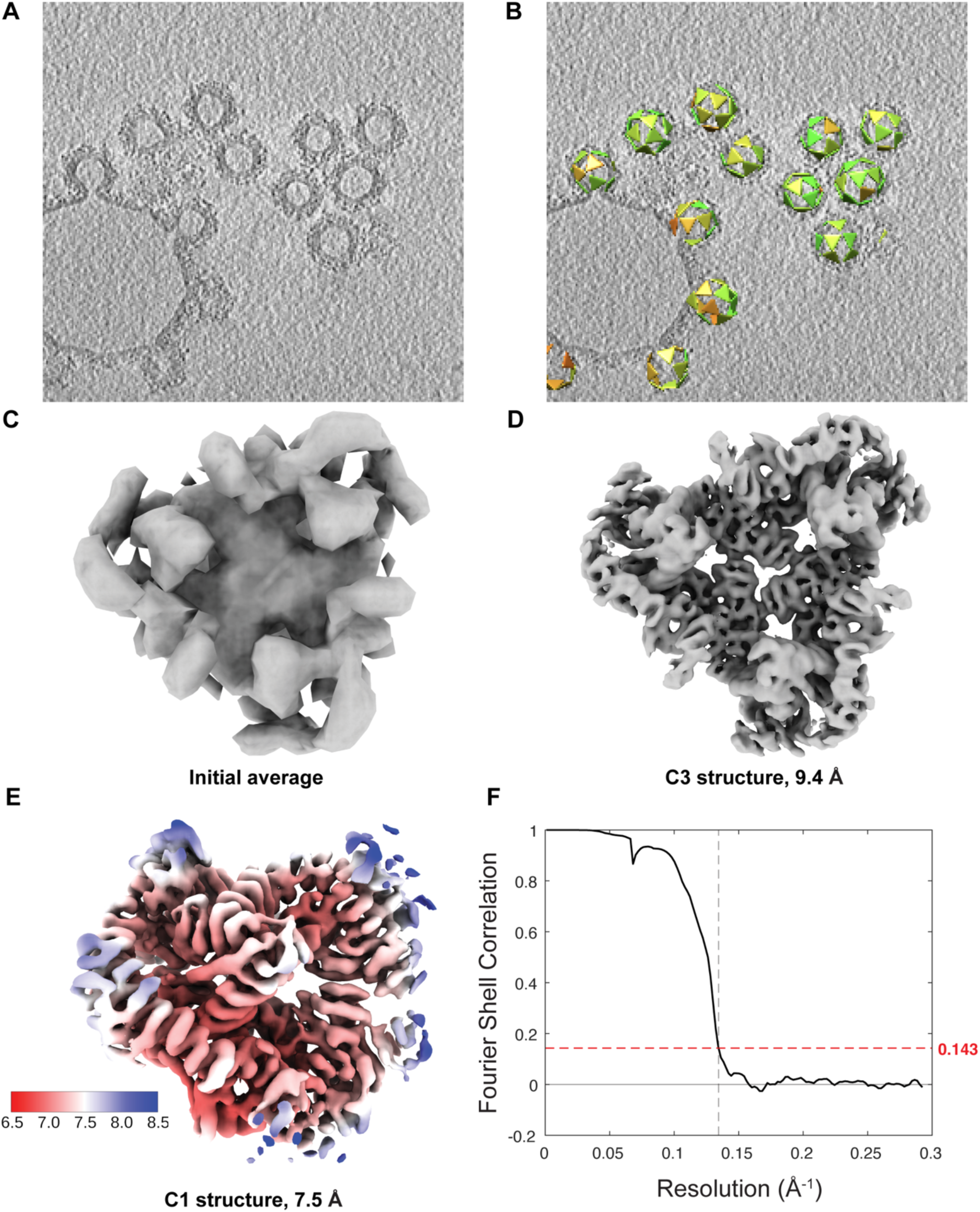
Subtomogram averaging of COPI-GOLPH3 complex. (A) Slice through a representative tomogram showing examples of COPI buds and vesicles. (B) Aligned COPI triad positions marked by triangles, superimposed on the tomogram slice. (C) initial average from SUBTOM picks after cleaning the particle list. (D) C3-symmetric COPI-GOLPH3 structure at 9.4 Å resolution. (E) C1 COPI-GOLPH3 structure obtained after symmetry expansion at 7.5 Å resolution. (F) Fourier shell correlation plot. See also table S1.

**Supplementary Figure 3.**
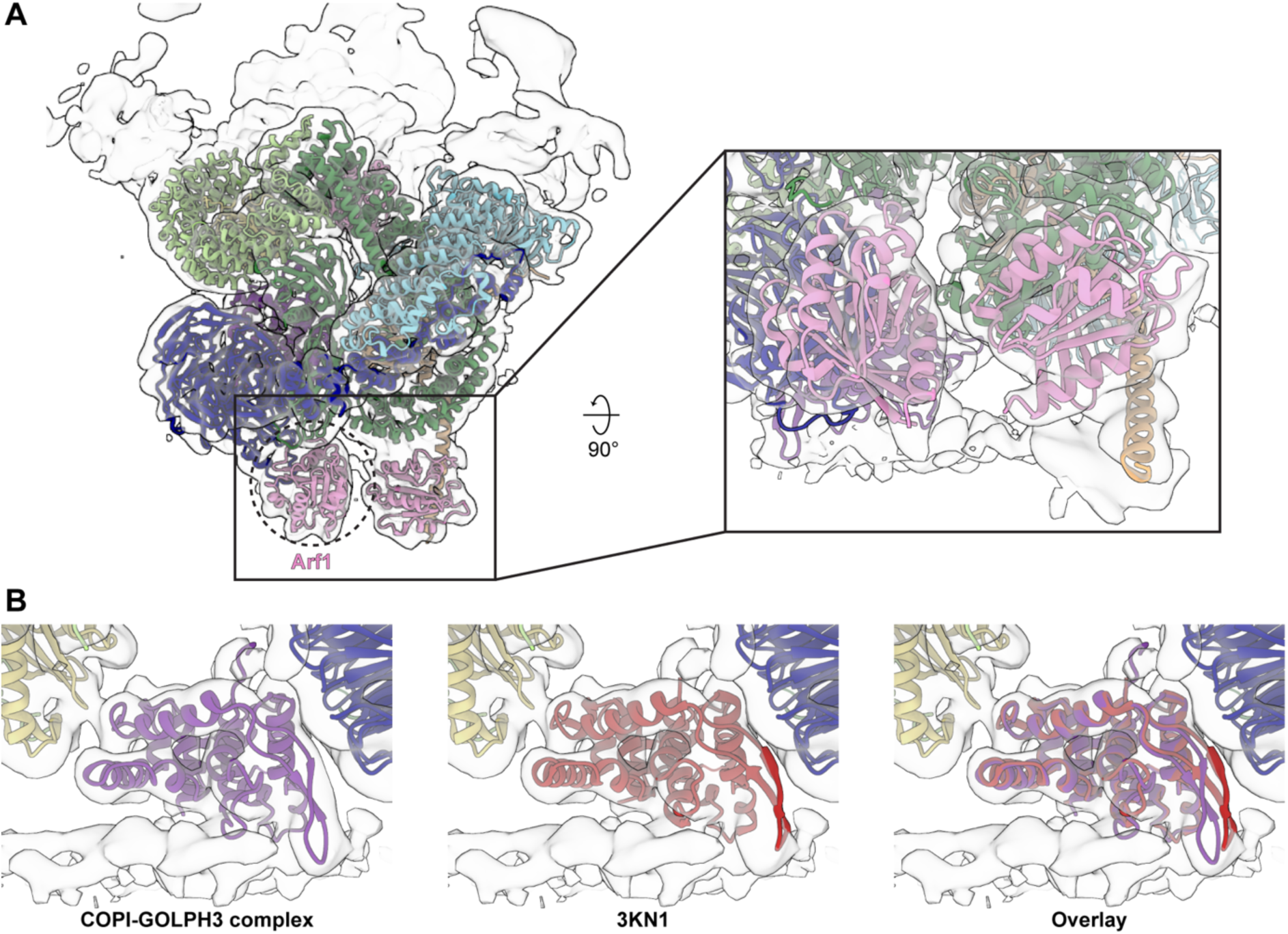
Comparisons of COPI-GOLPH3 complex to COPI and GOLPH3 structures. (A) The left panel shows the semi-transparent isosurface of the COPI leaf EM density map fitted with ribbon models of the COPI heptameric leaf containing GOLPH3 is shown from from the vesicle exterior looking down toward the membrane. Dotted circle marks density adjacent to the N-terminal beta-propellor of α-COP that was previously proposed to be constituted of a flexible loop in α-COP, but that can now be assigned as an additional copy of Arf1. The right panel shows a zoomed-in view, 90°-rotated to be perpendicular to the membrane to illustrate the fitting of the Arf1 ribbon model into the density. Components are coloured as follows: GOLPH3, purple, α-COP, dark blue; β-COP, dark green; β’-COP, light blue; δ-COP, orange; γ-COP, light green; ξ-COP, yellow, Arf1, pink. (B) Semi-transparent isosurface fitted with a ribbon model of the COPI-GOLPH3 leaf. The view is centered on GOLPH3 and is perpendicular to the membrane. For comparison, the center panel, shows the density fitted with the crystal structure of GOLPH3 (PDB: 3KN1, (*21*)). The right panel shows an overlay.

**Supplementary Figure 4.**
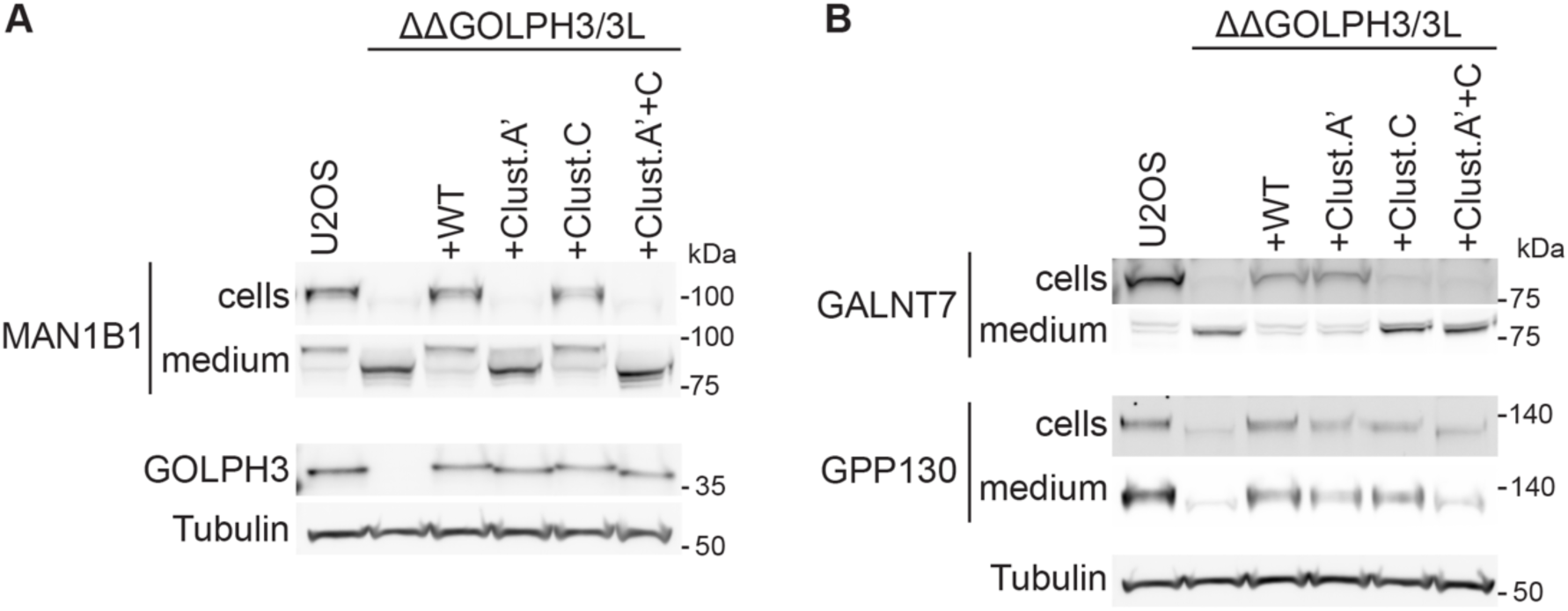
Golgi residents destabilised by loss of GOLPH3 are either clipped and secreted, or degraded intracellularly. (A) Immunoblots of whole-cell lysates and medium from either wild type U2OS cells or those or lacking both GOLPH3 and GOLPH3L with the latter transfected with either wild-type GOLPH3 (WT) or the indicated mutant forms. Blots were labelled for the Golgi enzyme MAN1B1, GOLPH3 or tubulin as a loading control. When MAN1B1 is not retained by GOLPH3 it is released into the medium in a clipped form. Such clipping has been seen with other Golgi residents when their localisation is perturbed (*31*). (B) Immunoblots of cells as in (A) but labelled for the Golgi residents GALNT7 and GPP130. When GALNT7 is not retained it is released into the medium in a clipped form. GPP130 does not appear in the medium at elevated levels when its cellular levels are reduced, consistent with previous reports that it is trafficked to the lysosome and degraded when not retained in the Golgi (*67*). We were unable to detect FAM3C in the medium and so could not determine its fate in the absence of GOLPH3-dependent retention.

**Supplementary Figure 5.**
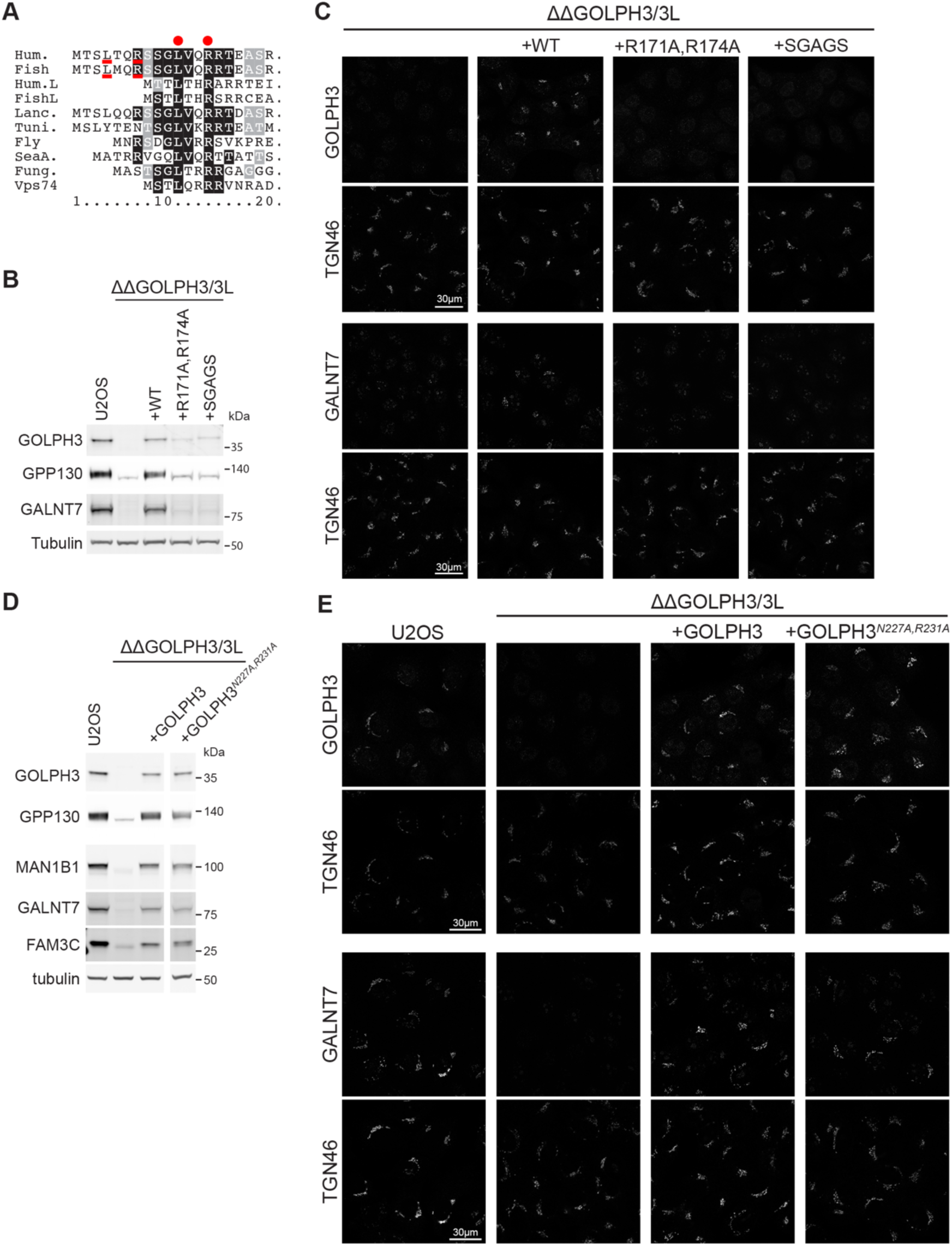
Mapping of parts of GOLPH3 required for function. (A) Alignment of the N-termini of GOLPH3 from the indicated species (Hum, human; Fish, coelacanth; Lanc, lancelet; Tuni, tunicate; Fly, *D. melanogaster*; SeaA, *Nematostella vectesis*; Fung, Aspergillus; Vps74, *S. cerevisiae*). Vertebrates also have GOLPH3L, as indicated. The conserved leucine and arginine at the ends of the region predicted to interact with coatomer are indicated with red dots. In vertebrate GOLPH3, this motif appears to be present twice, as indicated with underlining. (B) Immunoblot against GOLPH3 of whole cell lysates from U2OS cells deleted for GOLPH3 and GOLPH3L and expressing the indicated forms of GOLPH3. R171 and R174 are in the PI4P-binding site, and in SGAGS the hydrophobic hairpin residues _194_FLLFD_198_ are replaced with the eponymous sequence. (C) Confocal micrographs of cells as in (B), labelled for the Golgi marker TGN46 and either GOLPH3, or the GOLPH3-dependent Golgi resident GALNT7. (D) Immunoblots of whole cell lysates from U2OS ΔΔGOLPH3/3L expressing either nothing or GOLPH3 or GOLPH3 with mutations in residues near the possible interface with ζ-COP. Blots are labelled for GOLPH3 and the indicated Golgi residents or tubulin as a loading control, and the mutations do not appear to affect GOLPH3 function. (E) Confocal micrographs of cells as in (D), labelled as in (C).

**Supplementary Figure 6.**
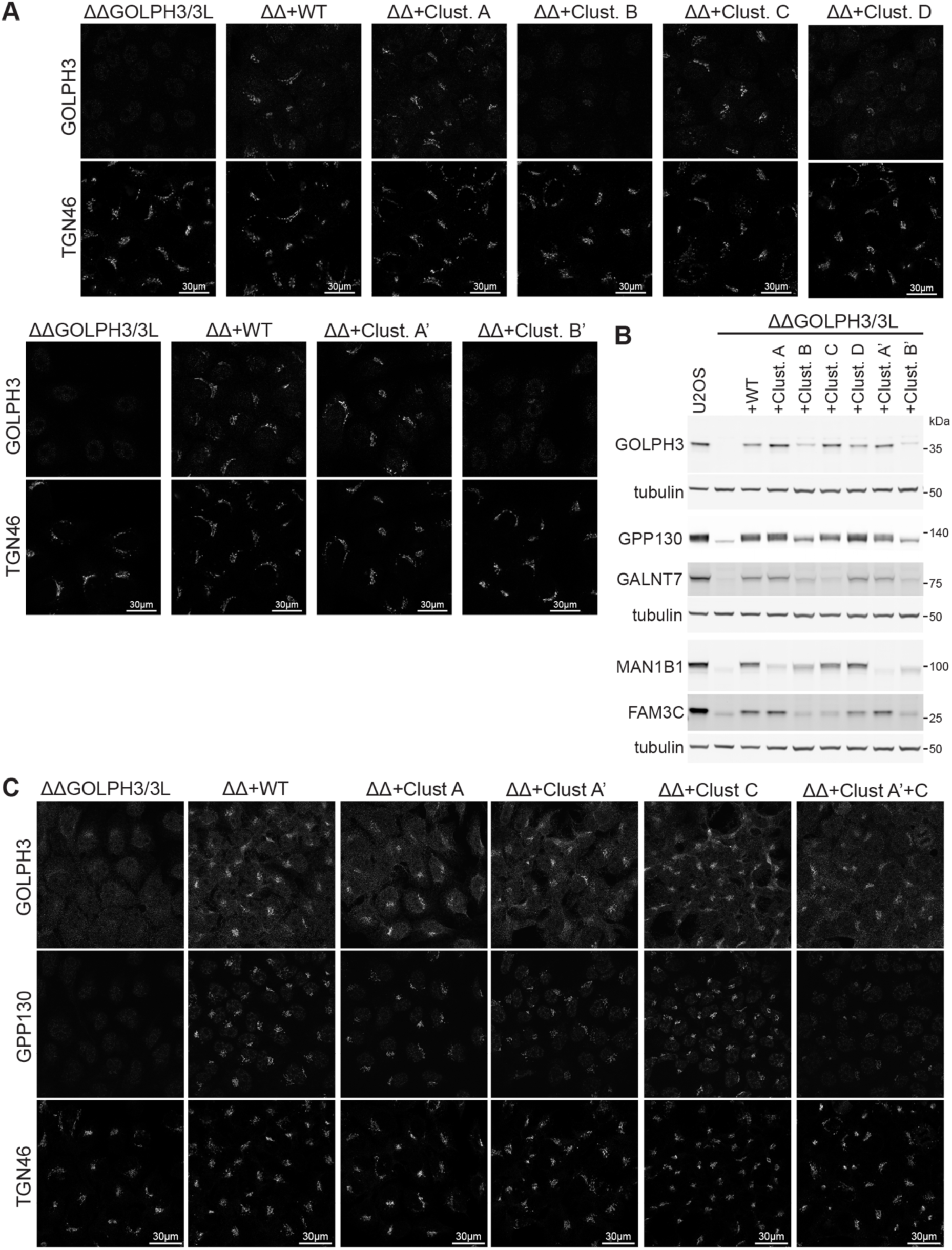
Effect on GOLPH3 function of mutating clusters of acidic residues. (A) Confocal micrographs of U2OS cells lacking GOLPH3 and GOLPH3L and transfected with either wild type GOLPH3 (WT), or forms with mutations to alanine of acidic residues in the indicated clusters. Cells were labelled for GOLPH3 and the Golgi marker TGN46. (B) Immunoblots of whole cell lysates from U2OS cells lacking GOLPH3 and GOLPH3L and transfected with either wild type GOLPH3 (WT), or forms with mutations to alanine of acidic residues in the indicated clusters. Blots were labelled for GOLPH3 and the indicated Golgi residents or tubulin as a loading control. Mutations in different clusters affect the ability of GOLPH3 to rescue the stability of different residents. (C) Confocal micrographs of U2OS cells lacking GOLPH3 and GOLPH3L and transfected with either wild type GOLPH3 (WT), or forms with mutations to alanine of acidic residues in the indicated clusters. Cells were labelled for GOLPH3, TGN46, and the GOLPH3-dependent Golgi resident GPP130.

**Supplementary Figure 7.**
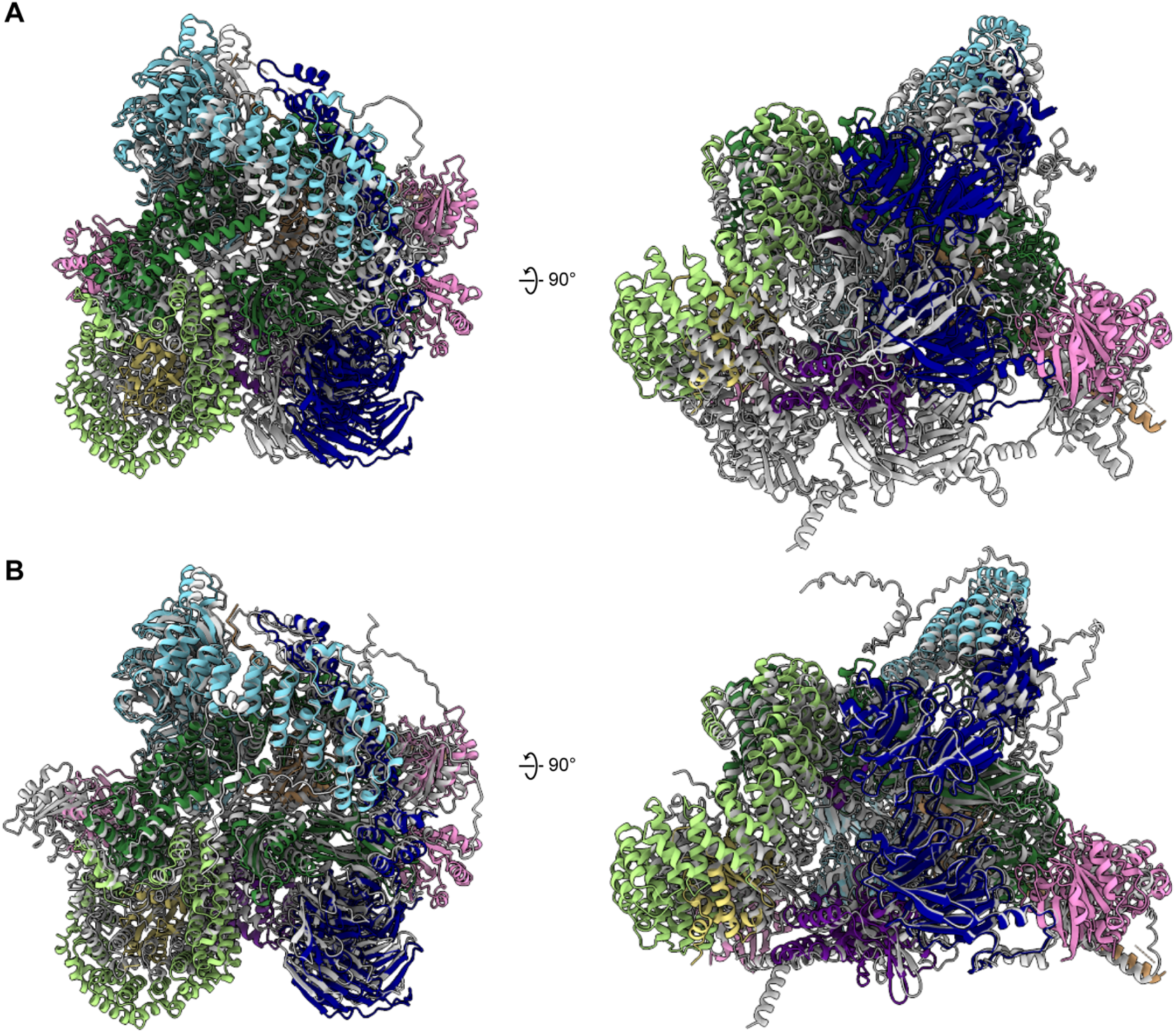
Comparisons of Alphafold 3 predictions of the structure of GOLPH3 bound to COPI to the final model. (A) The ribbon model of the COPI-GOLPH leaf (colored by component) aligned to a representative example of an Alphafold 3 prediction with the compressed conformation observed in approximately 70% of predictions (grey). (B) The ribbon model of the COPI-GOLPH leaf (colored by component) aligned to a representative example of an Alphafold 3 prediction with a conformation similar to that observed in the experimental density, observed in approximately 30% of predictions (grey). A prediction with the conformation B was used as the starting point for model building.

**Table S1.** CryoEM data acquisition and image processing.

| Sample | COPI leaf with *M. musculus* COPI. *H. sapiens* Arf1, *H. sapiens* GOLPH3 |
| --- | --- |
| **Data acquisition** |  |
| Microscope | FEI Titan Krios |
| Voltage (keV) | 300 |
| Energy-filter (eV) | 20 |
| Detector | Gatan BioQuantum K3 |
| Pixel size (Å) | 1.701 |
| Defocus range (microns) | -1.0 to -3.2 |
| Acquisition scheme | -60/60°, 3° |
| Total Dose (electrons/Å<sup>2</sup>) | 125 |
| Dose rate<br>(electrons/Å<sup>2</sup>/sec) | 8.56 |
| Frame number | 10 |
| Tomogram number | 206 |
| **Image processing** |  |
| Vesicles and buds | 4866 |
| Subtomograms | 108432 |
| Symmetry | C1 |
| Resolution at 0.143 FSC<br>(Å) | 7.5 |
| B factor | -600 |

**Table S2.**
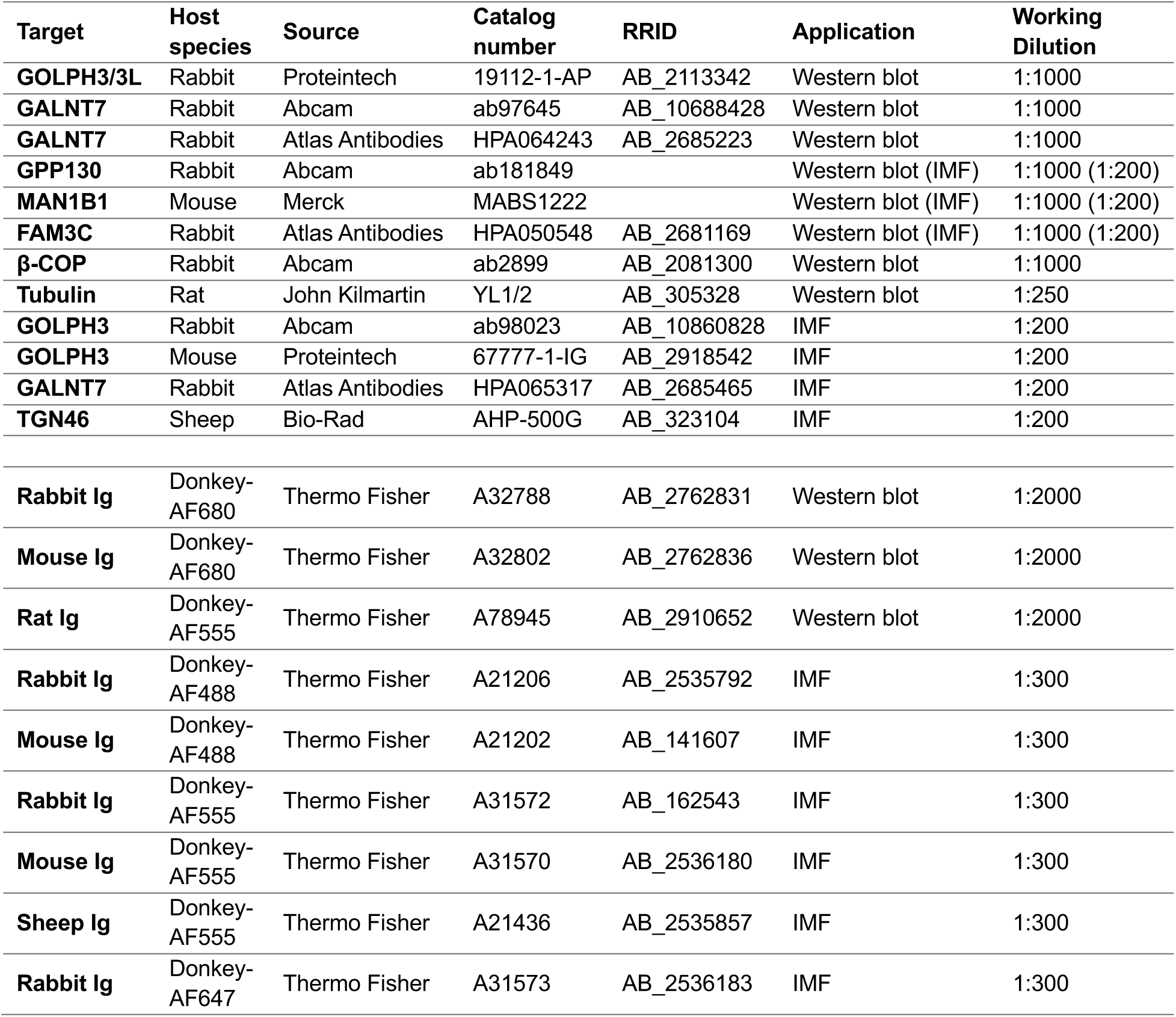
Primary and secondary antibodies used in this study. Fluorescent secondary antibodies for immunofluorescence (IMF) and western blotting were labelled with Alexa Fluor (AF) dyes.

| Target | Host species | Source | Catalog number | RRID | Application | Working Dilution |
| --- | --- | --- | --- | --- | --- | --- |
| **GOLPH3/3L** | Rabbit | Proteintech | 19112-1-AP | AB_2113342 | Western blot | 1:1000 |
| **GALNT7** | Rabbit | Abcam | ab97645 | AB_10688428 | Western blot | 1:1000 |
| **GALNT7** | Rabbit | Atlas Antibodies | HPA064243 | AB_2685223 | Western blot | 1:1000 |
| **GPP130** | Rabbit | Abcam | ab181849 |  | Western blot (IMF) | 1:1000 (1:200) |
| **MAN1B1** | Mouse | Merck | MABS1222 |  | Western blot (IMF) | 1:1000 (1:200) |
| **FAM3C** | Rabbit | Atlas Antibodies | HPA050548 | AB_2681169 | Western blot (IMF) | 1:1000 (1:200) |
| **β-COP** | Rabbit | Abcam | ab2899 | AB_2081300 | Western blot | 1:1000 |
| **Tubulin** | Rat | John Kilmartin | YL1/2 | AB_305328 | Western blot | 1:250 |
| **GOLPH3** | Rabbit | Abcam | ab98023 | AB_10860828 | IMF | 1:200 |
| **GOLPH3** | Mouse | Proteintech | 67777-1-IG | AB_2918542 | IMF | 1:200 |
| **GALNT7** | Rabbit | Atlas Antibodies | HPA065317 | AB_2685465 | IMF | 1:200 |
| **TGN46** | Sheep | Bio-Rad | AHP-500G | AB_323104 | IMF | 1:200 |
| **Rabbit Ig** | Donkey-AF680 | Thermo Fisher | A32788 | AB_2762831 | Western blot | 1:2000 |
| **Mouse Ig** | Donkey-AF680 | Thermo Fisher | A32802 | AB_2762836 | Western blot | 1:2000 |
| **Rat Ig** | Donkey-AF555 | Thermo Fisher | A78945 | AB_2910652 | Western blot | 1:2000 |
| **Rabbit Ig** | Donkey-AF488 | Thermo Fisher | A21206 | AB_2535792 | IMF | 1:300 |
| **Mouse Ig** | Donkey-AF488 | Thermo Fisher | A21202 | AB_141607 | IMF | 1:300 |
| **Rabbit Ig** | Donkey-AF555 | Thermo Fisher | A31572 | AB_162543 | IMF | 1:300 |
| **Mouse Ig** | Donkey-AF555 | Thermo Fisher | A31570 | AB_2536180 | IMF | 1:300 |
| **Sheep Ig** | Donkey-AF555 | Thermo Fisher | A21436 | AB_2535857 | IMF | 1:300 |
| **Rabbit Ig** | Donkey-AF647 | Thermo Fisher | A31573 | AB_2536183 | IMF | 1:300 |

## References

1. B. S. Glick, A. Nakano, Membrane traffic within the Golgi apparatus. Annu Rev Cell Dev Biol 25, 113–132 (2009).

2. M. G. Farquhar, G. E. Palade, The Golgi apparatus: 100 years of progress and controversy. Trends in Cell Biology 8, 2–10 (1998).

3. K. W. Moremen, M. Tiemeyer, A. V. Nairn, Vertebrate protein glycosylation: diversity, synthesis and function. Nat Rev Mol Cell Biol 13, 448–462 (2012).

4. K. T. Schjoldager, Y. Narimatsu, H. J. Joshi, H. Clausen, Global view of human protein glycosylation pathways and functions. Nat Rev Mol Cell Biol 21, 729– 749 (2020).

5. J. Béthune, F. T. Wieland, Assembly of COPI and COPII Vesicular Coat Proteins on Membranes. Annu. Rev. Biophys. 47, 63–83 (2018).

6. N. Gomez-Navarro, E. Miller, Protein sorting at the ER-Golgi interface. J Cell Biol 215, 769–778 (2016).

7. B. S. Glick, A. Luini, Models for Golgi traffic: a critical assessment. Cold Spring Harb Perspect Biol 3, a005215 (2011).

8. S. Emr, B. S. Glick, A. D. Linstedt, J. Lippincott-Schwartz, A. Luini, V. Malhotra, B. J. Marsh, A. Nakano, S. R. Pfeffer, C. Rabouille, J. E. Rothman, G. Warren, F. T. Wieland, Journeys through the Golgi-taking stock in a new era. J. Cell Biol. 187, 449–453 (2009).

9. Y. S. Bykov, M. Schaffer, S. O. Dodonova, S. Albert, J. M. Plitzko, W. Baumeister, B. D. Engel, J. A. Briggs, The structure of the COPI coat determined within the cell. eLife 6, e32493 (2017).

10. R. J. Taylor, G. Tagiltsev, J. A. G. Briggs, The structure of COPI vesicles and regulation of vesicle turnover. FEBS Letters, 1873-3468.14560 (2022).

11. E. C. Arakel, B. Schwappach, Formation of COPI-coated vesicles at a glance. J Cell Sci 131, jcs209890 (2018).

12. L. P. Jackson, M. Lewis, H. M. Kent, M. A. Edeling, P. R. Evans, R. Duden, D. J. Owen, Molecular basis for recognition of dilysine trafficking motifs by COPI. Dev Cell 23, 1255–1262 (2012).

13. W. Ma, J. Goldberg, Rules for the recognition of dilysine retrieval motifs by coatomer. EMBO J 32, 926–937 (2013).

14. L. G. Welch, S. Munro, A tale of short tails, through thick and thin: investigating the sorting mechanisms of Golgi enzymes. FEBS Lett 593, 2452–2465 (2019).

15. P. Lujan, F. Campelo, Should I stay or should I go? Golgi membrane spatial organization for protein sorting and retention. Arch Biochem Biophys 707, 108921 (2021).

16. L. Tu, W. C. S. Tai, L. Chen, D. K. Banfield, Signal-mediated dynamic retention of glycosyltransferases in the Golgi. Science 321, 404–407 (2008).

17. K. R. Schmitz, J. Liu, S. Li, T. G. Setty, C. S. Wood, C. G. Burd, K. M. Ferguson, Golgi localization of glycosyltransferases requires a Vps74p oligomer. Dev Cell 14, 523–534 (2008).

18. E. S. P. Eckert, I. Reckmann, A. Hellwig, S. Röhling, A. El-Battari, F. T. Wieland, V. Popoff, Golgi phosphoprotein 3 triggers signal-mediated incorporation of glycosyltransferases into coatomer-coated (COPI) vesicles. J Biol Chem 289, 31319–31329 (2014).

19. L. G. Welch, S.-Y. Peak-Chew, F. Begum, T. J. Stevens, S. Munro, GOLPH3 and GOLPH3L are broad-spectrum COPI adaptors for sorting into intra-Golgi transport vesicles. J Cell Biol 220, e202106115 (2021).

20. L. Tu, L. Chen, D. K. Banfield, A conserved N-terminal arginine-motif in GOLPH3-family proteins mediates binding to coatomer. Traffic 13, 1496–1507 (2012).

21. C. S. Wood, K. R. Schmitz, N. J. Bessman, T. G. Setty, K. M. Ferguson, C. G. Burd, PtdIns4P recognition by Vps74/GOLPH3 links PtdIns 4-kinase signaling to retrograde Golgi trafficking. J Cell Biol 187, 967–975 (2009).

22. S. Sechi, A. Frappaolo, A. Karimpour-Ghahnavieh, R. Piergentili, M. G. Giansanti, Oncogenic Roles of GOLPH3 in the Physiopathology of Cancer. Int J Mol Sci 21 (2020).

23. S. S. Pinho, C. A. Reis, Glycosylation in cancer: mechanisms and clinical implications. Nat Rev Cancer 15, 540–555 (2015).

24. B. K. Brauer, Z. Chen, F. Beirow, J. Li, D. Meisinger, E. Capriotti, M. Schweizer, L. Wagner, J. Wienberg, L. Hobohm, L. Blume, W. Qiao, Y. Narimatsu, J. E. Carette, H. Clausen, D. Winter, T. Braulke, S. Jabs, M. Voss, GOLPH3 and GOLPH3L maintain Golgi localization of LYSET and a functional mannose 6-phosphate transport pathway. EMBO J 43, 6264–6290 (2024).

25. R. Rizzo, D. Russo, K. Kurokawa, P. Sahu, B. Lombardi, D. Supino, M. A. Zhukovsky, A. Vocat, P. Pothukuchi, V. Kunnathully, L. Capolupo, G. Boncompain, C. Vitagliano, F. Zito Marino, G. Aquino, D. Montariello, P. Henklein, L. Mandrich, G. Botti, H. Clausen, U. Mandel, T. Yamaji, K. Hanada, A. Budillon, F. Perez, S. Parashuraman, Y. A. Hannun, A. Nakano, D. Corda, G. D’Angelo, A. Luini, Golgi maturation-dependent glycoenzyme recycling controls glycosphingolipid biosynthesis and cell growth via GOLPH3. EMBO J 40, e107238 (2021).

26. H. J. F. Maccioni, R. Quiroga, W. Spessott, Organization of the synthesis of glycolipid oligosaccharides in the Golgi complex. FEBS Lett 585, 1691–1698 (2011).

27. S. O. Dodonova, P. Aderhold, J. Kopp, I. Ganeva, S. Röhling, W. J. H. Hagen, I. Sinning, F. Wieland, J. A. G. Briggs, 9Å structure of the COPI coat reveals that the Arf1 GTPase occupies two contrasting molecular environments. eLife 6, e26691 (2017).

28. M. Faini, S. Prinz, R. Beck, M. Schorb, J. D. Riches, K. Bacia, B. Brügger, F. T. Wieland, J. A. G. Briggs, The Structures of COPI-Coated Vesicles Reveal Alternate Coatomer Conformations and Interactions. Science 336, 1451–1454 (2012).

29. S. O. Dodonova, P. Diestelkoetter-Bachert, A. Von Appen, W. J. H. Hagen, R. Beck, M. Beck, F. Wieland, J. A. G. Briggs, A structure of the COPI coat and the role of coat proteins in membrane vesicle assembly. Science 349, 195–198 (2015).

30. H. C. Dippold, M. M. Ng, S. E. Farber-Katz, S.-K. Lee, M. L. Kerr, M. C. Peterman, R. Sim, P. A. Wiharto, K. A. Galbraith, S. Madhavarapu, G. J. Fuchs, T. Meerloo, M. G. Farquhar, H. Zhou, S. J. Field, GOLPH3 bridges phosphatidylinositol-4-phosphate and actomyosin to stretch and shape the Golgi to promote budding. Cell 139, 337–351 (2009).

31. M. Voss, Proteolytic cleavage of Golgi glycosyltransferases by SPPL3 and other proteases and its implications for cellular glycosylation. Biochimica et Biophysica Acta (BBA) - General Subjects 1868, 130668 (2024).

32. Y. Cai, Y. Deng, F. Horenkamp, K. M. Reinisch, C. G. Burd, Sac1-Vps74 structure reveals a mechanism to terminate phosphoinositide signaling in the Golgi apparatus. J Cell Biol 206, 485–491 (2014).

33. M. G. Farquhar, G. E. Palade, The Golgi apparatus (complex)-(1954-1981)-from artifact to center stage. J. Cell Biol. 91, 77s–103s (1981).

34. W. E. Balch, W. G. Dunphy, W. A. Braell, J. E. Rothman, Reconstitution of the transport of protein between successive compartments of the golgi measured by the coupled incorporation of N-acetylglucosamine. Cell 39, 405–416 (1984).

35. M. G. Waters, T. Serafini, J. E. Rothman, “Coatomer”: a cytosolic protein complex containing subunits of non-clathrin-coated Golgi transport vesicles. Nature 349, 248–251 (1991).

36. F. Letourneur, E. C. Gaynor, S. Hennecke, C. Démollière, R. Duden, S. D. Emr, H. Riezman, P. Cosson, Coatomer is essential for retrieval of dilysine-tagged proteins to the endoplasmic reticulum. Cell 79, 1199–1207 (1994).

37. A. Pantazopoulou, B. S. Glick, A kinetic view of membrane traffic pathways can transcend the classical view of Golgi compartments. Front Cell Dev Biol 7, 153 (2019).

38. A. Nakano, A. Luini, Passage through the Golgi. Curr Opin Cell Biol 22, 471– 478 (2010).

39. M. H. Dunlop, A. M. Ernst, L. K. Schroeder, D. K. Toomre, G. Lavieu, J. E. Rothman, Land-locked mammalian Golgi reveals cargo transport between stable cisternae. Nat Commun 8, 432 (2017).

40. 40. L. Orci, M. Ravazzola, A. Volchuk, T. Engel, M. Gmachl, M. Amherdt, A. Perrelet, T. H. Söllner, J. E. Rothman, Anterograde flow of cargo across the Golgi stack potentially mediated via bidirectional “percolating” COPI vesicles. Proc. Natl. Acad. Sci. U.S.A. 97, 10400–10405 (2000).

41. T. Szul, E. Sztul, COPII and COPI Traffic at the ER-Golgi Interface. Physiology 26, 348–364 (2011).

42. D. J. Stephens, N. Lin-Marq, A. Pagano, R. Pepperkok, J.-P. Paccaud, COPI-coated ER-to-Golgi transport complexes segregate from COPII in close proximity to ER exit sites. Journal of Cell Science 113, 2177–2185 (2000).

43. T. Baba, A. Alvarez-Prats, Y. J. Kim, D. Abebe, S. Wilson, Z. Aldworth, M. A. Stopfer, J. Heuser, T. Balla, Myelination of peripheral nerves is controlled by PI4KB through regulation of Schwann cell Golgi function. Proc. Natl. Acad. Sci. U.S.A. 117, 28102–28113 (2020).

44. Y. Liu, M. Boukhelifa, E. Tribble, E. Morin-Kensicki, A. Uetrecht, J. E. Bear, V. A. Bankaitis, The Sac1 Phosphoinositide Phosphatase Regulates Golgi Membrane Morphology and Mitotic Spindle Organization in Mammals. MBoC 19, 3080–3096 (2008)

45. M. Ishida, ARMH3 is an ARL5 effector that promotes PI4KB-catalyzed PI4P synthesis at the trans-Golgi network. Nature Communications (2024) 15:10168.

46. M. Y. Hein, D. Peng, V. Todorova, F. McCarthy, K. Kim, C. Liu, L. Savy, C. Januel, R. Baltazar-Nunez, M. Sekhar, S. Vaid, S. Bax, M. Vangipuram, J. Burgess, L. Njoya, E. Wang, I. E. Ivanov, J. R. Byrum, S. Pradeep, C. G. Gonzalez, Y. Aniseia, J. S. Creery, A. H. McMorrow, S. Sunshine, S. Yeung-Levy, B. C. DeFelice, S. B. Mehta, D. N. Itzhak, J. E. Elias, M. D. Leonetti, Global organelle profiling reveals subcellular localization and remodeling at proteome scale. Cell 188, 1137–1155.e20 (2025).

47. H. C. Tie, D. Mahajan, L. Lu, Visualizing intra-Golgi localization and transport by side-averaging Golgi ministacks. Journal of Cell Biology 221, e202109114 (2022).

48. L. Liu, N. Watanabe, H. Akatsu, M. Nishimura, Neuronal expression of ILEI/FAM3C and its reduction in Alzheimer’s disease. Neuroscience 330, 236– 246 (2016).

49. N. H. Cho, K. C. Cheveralls, A.-D. Brunner, K. Kim, A. C. Michaelis, P. Raghavan, H. Kobayashi, L. Savy, J. Y. Li, H. Canaj, J. Y. S. Kim, E. M. Stewart, C. Gnann, F. McCarthy, J. P. Cabrera, R. M. Brunetti, B. B. Chhun, G. Dingle, M. Y. Hein, B. Huang, S. B. Mehta, J. S. Weissman, R. Gómez-Sjöberg, D. N. Itzhak, L. A. Royer, M. Mann, M. D. Leonetti, OpenCell: Endogenous tagging for the cartography of human cellular organization. Science 375, eabi6983 (2022).

50. M. C. Sahlmüller, J. R. P. M. Strating, R. Beck, P. Eckert, V. Popoff, M. Haag, A. Hellwig, I. Berger, B. Brügger, F. T. Wieland, Recombinant heptameric coatomer complexes: novel tools to study isoform-specific functions. Traffic 12, 682–692 (2011).

51. P. A. Randazzo, O. Weiss, R. A. Kahn, “[34] Preparation of recombinant ADP-ribosylation factor” in Methods in Enzymology, J. E. Rothman, Ed. (Academic Press, 1992)vol. 219 *of Reconstitution of Intracellular* Transport, pp. 362–369.

52. E. Mossessova, J. M. Gulbis, J. Goldberg, Structure of the guanine nucleotide exchange factor Sec7 domain of human arno and analysis of the interaction with ARF GTPase. Cell 92, 415–423 (1998).

53. A. Spang, K. Matsuoka, S. Hamamoto, R. Schekman, L. Orci, Coatomer, Arf1p, and nucleotide are required to bud coat protein complex I-coated vesicles from large synthetic liposomes. Proc. Natl. Acad. Sci. U.S.A. 95, 11199–11204 (1998).

54. W. J. H. Hagen, W. Wan, J. A. G. Briggs, Implementation of a cryo-electron tomography tilt-scheme optimized for high resolution subtomogram averaging. Journal of Structural Biology 197, 191–198 (2017).

55. D. N. Mastronarde, Automated electron microscope tomography using robust prediction of specimen movements. Journal of Structural Biology 152, 36–51 (2005).

56. J. R. Kremer, D. N. Mastronarde, J. R. McIntosh, Computer Visualization of Three-Dimensional Image Data Using IMOD. Journal of Structural Biology 116, 71–76 (1996).

57. T. Grant, N. Grigorieff, Automatic estimation and correction of anisotropic magnification distortion in electron microscopes. Journal of Structural Biology 192, 204–208 (2015).

58. Q. Xiong, M. K. Morphew, C. L. Schwartz, A. H. Hoenger, D. N. Mastronarde, CTF determination and correction for low dose tomographic tilt series. Journal of Structural Biology 168, 378–387 (2009).

59. J. G. Galaz-Montoya, J. Flanagan, M. F. Schmid, S. J. Ludtke, Single particle tomography in EMAN2. Journal of Structural Biology 190, 279–290 (2015).

60. B. Turoňová, F. K. M. Schur, W. Wan, J. A. G. Briggs, Efficient 3D-CTF correction for cryo-electron tomography using NovaCTF improves subtomogram averaging resolution to 3.4 Å. Journal of Structural Biology 199, 187–195 (2017).

61. E. F. Pettersen, T. D. Goddard, C. C. Huang, G. S. Couch, D. M. Greenblatt, E. C. Meng, T. E. Ferrin, UCSF Chimera--a visualization system for exploratory research and analysis. J Comput Chem 25, 1605–1612 (2004).

62. K. Qu, Z. Ke, V. Zila, M. Anders-Össwein, B. Glass, F. Mücksch, R. Müller, C. Schultz, B. Müller, H.-G. Kraüsslich, J. A. G. Briggs, Maturation of the matrix and viral membrane of HIV-1. Science 373, 700–704 (2021).

63. D. Tegunov, P. Cramer, Real-time cryo-electron microscopy data preprocessing with Warp. Nat Methods 16, 1146–1152 (2019).

64. J. Zivanov, T. Nakane, B. O. Forsberg, D. Kimanius, W. J. Hagen, E. Lindahl, S. H. Scheres, New tools for automated high-resolution cryo-EM structure determination in RELION-3. eLife 7, 163 (2018).

65. D. Tegunov, L. Xue, C. Dienemann, P. Cramer, J. Mahamid, Multi-particle cryo-EM refinement with M visualizes ribosome-antibiotic complex at 3.5 Å in cells. Nat Methods 18, 186–193 (2021).

66. J. Abramson, J. Adler, J. Dunger, R. Evans, T. Green, A. Pritzel, O. Ronneberger, L. Willmore, A. J. Ballard, J. Bambrick, S. W. Bodenstein, D. A. Evans, C.-C. Hung, M. O’Neill, D. Reiman, K. Tunyasuvunakool, Z. Wu, A. Žemgulytė, E. Arvaniti, C. Beattie, O. Bertolli, A. Bridgland, A. Cherepanov, M. Congreve, A. I. Cowen-Rivers, A. Cowie, M. Figurnov, F. B. Fuchs, H. Gladman, R. Jain, Y. A. Khan, C. M. R. Low, K. Perlin, A. Potapenko, P. Savy, S. Singh, A. Stecula, A. Thillaisundaram, C. Tong, S. Yakneen, E. D. Zhong, M. Zielinski, A. Žídek, V. Bapst, P. Kohli, M. Jaderberg, D. Hassabis, J. M. Jumper, Accurate structure prediction of biomolecular interactions with AlphaFold 3. Nature 630, 493–500 (2024).

67. S. Mukhopadhyay, C. Bachert, D. R. Smith, A. D. Linstedt, Manganese-induced Trafficking and Turnover of the *cis* -Golgi Glycoprotein GPP130. MBoC 21, 1282–1292 (2010).

